# p53 is a selective barrier to chromosome loss in cancer

**DOI:** 10.1101/2025.07.04.663162

**Authors:** Joana F. Marques, Marco António Dias Louro, Michael Schubert, Tom Winkler, Guy Wolf-Dankovich, Teresa Davoli, Uri Ben-David, Geert J.P.L. Kops

**Affiliations:** Hubrecht Institute, Royal Netherlands Academy of Arts and Sciences (KNAW), Uppsalalaan 8, 3584CT Utrecht, the Netherlands; University Medical Center Utrecht, Heidelberglaan 100, 3584CX Utrecht, the Netherlands; Oncode Institute, Jaarbeursplein 6, 3521AL Utrecht, the Netherlands; Institute of Bioinformatics, Medical University of Innsbruck, Innrain 80, 6020 Innsbruck, Austria; Department of Human Genetics and Computational Medicine, Gray Faculty of Medical and Health Sciences, Tel Aviv University, Tel Aviv, Israel; Institute for Systems Genetics and Department of Biochemistry and Molecular Pharmacology, NYU Grossman School of Medicine, New York, NY, USA

## Abstract

Aneuploidy and *TP53* mutations are among the most frequent genetic alterations in cancer. p53 inactivation is widely considered a key enabling event for aneuploidy emergence. However, whether p53 protects against specific types of aneuploidy, and whether this is universal across cancer types, remains unclear. By analyzing aneuploidy features across cancer types in the TCGA database, we show that p53 inactivation is neither sufficient nor necessary for cancer aneuploidy. While p53-deficient tumors show higher pan-cancer aneuploidy, within individual cancer types p53-proficient and -deficient tumors often have comparable aneuploidy levels, and p53-proficient cancers can be highly aneuploid. Instead, p53 deficiency is associated with a higher frequency of chromosome losses. Single-cell sequencing of non-transformed cells after induced chromosomal instability reveals preferential p53 pathway activation upon chromosome loss but not gain. Prolonged culture selectively enriches for monosomies when p53 is inactivated. p53 thus acts as a context-dependent safeguard against aneuploidy, particularly against chromosome losses.

## Introduction

Aneuploidy, here defined as copy-number alterations (CNAs) of whole chromosomes or chromosome arms, is the most common genetic alteration in cancer, affecting 75-90% of tumors (*1-4*). Aneuploidy contributes to tumor initiation, metastasis, and therapy resistance (*5-7*), yet it is detrimental to healthy cells (*8-11*). The most common genetic alteration in cancer next to aneuploidy is inactivating mutations in the *TP53* tumor suppressor gene, affecting ∼41% of tumors (*1, 3, 12, 13*). Their strong co-occurrence in cancer and the well-described role of p53 in maintaining genome stability suggests a causal association between p53 inactivation and cancer aneuploidy emergence. This was supported by pan-cancer genome analyses (*3, 13-16*) and experimental studies of aneuploidy tolerance upon p53 inactivation in non-transformed cells (*17-20*). Nevertheless, aneuploidy occurs more frequently in cancer than do mutations in *TP53*, and recent experimental studies using human and murine intestinal and mammary organoid cultures have challenged the causal link between the two (*21, 22*). It is thus unclear whether p53 inactivation is important for cancer aneuploidy *per se* or whether it is, for example, more specifically relevant for certain cancer types and/or for emergence of particular features of aneuploidy such as degree, type (arm, whole chromosome or both) or direction (gains vs losses) (*18*). To better understand the contribution of p53 to protection against aneuploidy, we here investigated cancer genome data for associations between p53 status and various aneuploidy features, and validated findings experimentally in non-transformed human cell lines.

## Results

### The association of aneuploidy with p53 deficiency is cancer type-specific

To examine the association of aneuploidy and p53 status in cancer, we explored TCGA data from 8539 tumors across 31 cancer types for various associations between chromosomal copy number, p53 status, and ploidy (Fig. 1A, Suppl. Table 1A, and see methods for details on CNA, p53 status and ploidy calling). Tumor samples were considered aneuploidy if containing at least one arm or whole chromosome CNA. This resulted in the classification of 1144 samples as euploid and 7395 samples as aneuploid, of which 40.3% (2978) were designated as having undergone a whole genome doubling (WGD, see Suppl. Fig. 1, Suppl. Table 1B). For the three categories (euploid, aneuploid, aneuploid with WGD), we designated samples as proficient (+) or deficient (–) for p53 function in the following manner: first, all samples with predicted inactivating mutations in the *TP53* gene (see methods) were classified as p53 deficient; second, we used *TP53*-loss phenocopy data from transcriptomic analyses to assess which samples with an intact *TP53* gene were likely to be p53 deficient by means other than *TP53* gene mutations, including mutations in p53 pathway components (*23*). 60% of non-WGD samples (2237 tumors) could be assessed for *TP53*-loss phenocopy, which resulted in the re-classification of 12% (299 of 2488) of *TP53* wild type samples as being p53 deficient (Suppl. Fig. 2).

**Figure 1.**
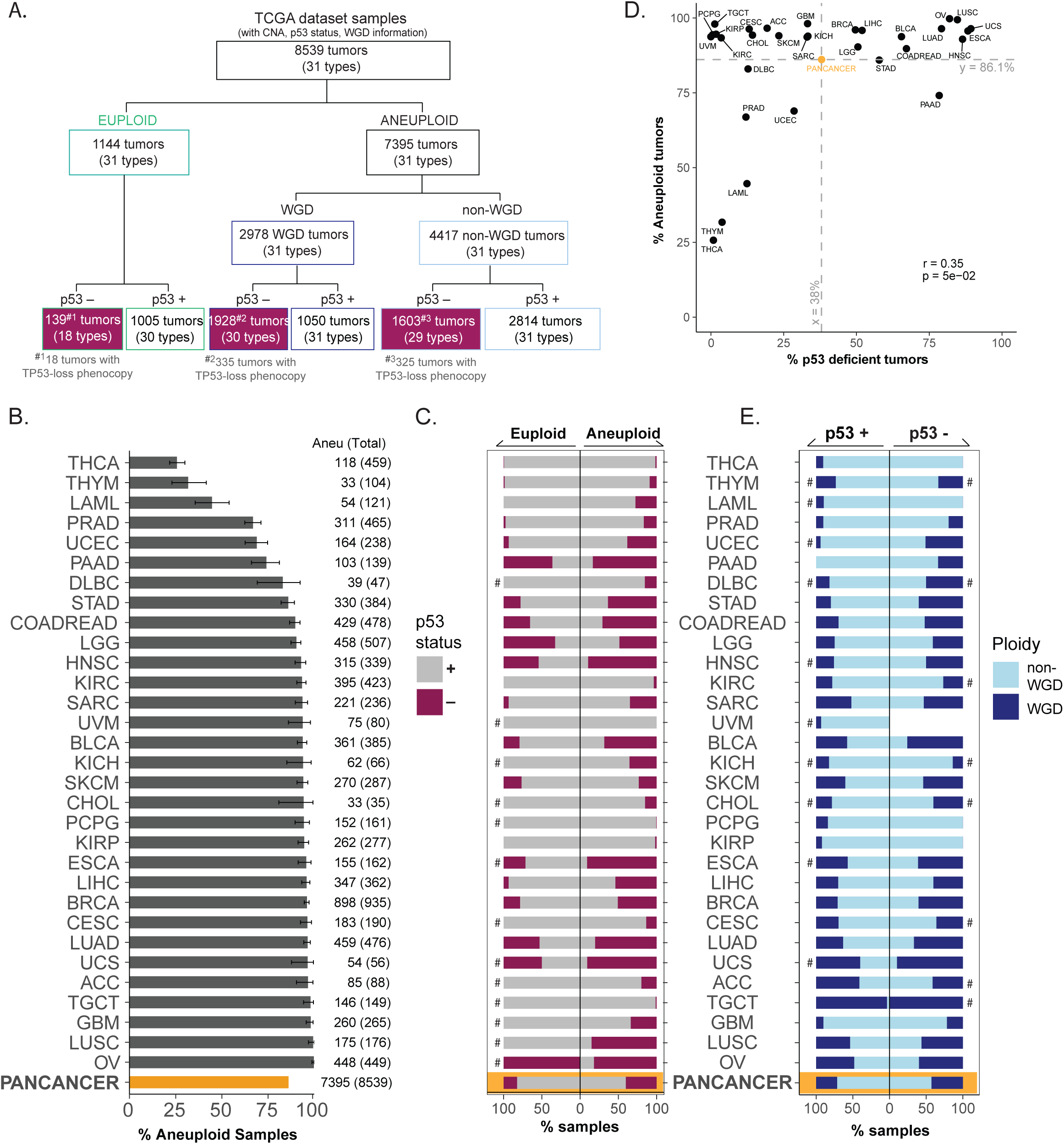
The association of aneuploidy with p53 deficiency is cancer type-specific. A. Subcategorization of TCGA samples based on aneuploidy (euploid and aneuploid), WGD (non-WGD and WGD), and p53 status (proficient (+) or deficient (–)). Number of samples with p53 loss phenocopy included are indicated below each respective category (grey text). B. Prevalence of aneuploidy across cancer types. The yellow bar depicts pan-cancer aneuploidy prevalence. Aneu (Total) - represents the number of aneuploid samples (total number of samples). Error bars indicate 95% confidence interval. Acute Myeloid Leukemia (LAML), Adrenocortical carcinoma (ACC), Bladder Urothelial Carcinoma (BLCA), Breast carcinoma (BRCA), Cervical squamous cell carcinoma and endocervical adenocarcinoma (CESC), Cholangiocarcinoma (CHOL), Colorectal adenocarcinoma (COADREAD), Lymphoid Neoplasm Diffuse Large B-cell Lymphoma (DLBC), Esophageal carcinoma (ESCA), Glioblastoma multiforme (GBM), Head and Neck squamous cell carcinoma (HNSC), Kidney Chromophobe (KICH), Kidney renal clear cell carcinoma (KIRC), Kidney renal papillary cell carcinoma (KIRP), Brain Lower Grade Glioma (LGG), Liver hepatocellular carcinoma (LIHC), Lung adenocarcinoma (LUAD), Lung squamous cell carcinoma (LUSC), Ovarian serous cystadenocarcinoma (OV), Pancreatic adenocarcinoma (PAAD), Pheochromocytoma and Paraganglioma (PCPG), Prostate adenocarcinoma (PRAD), Sarcoma (SARC), Skin Cutaneous Melanoma (SKCM), Stomach adenocarcinoma (STAD), Testicular Germ Cell Tumors (TGCT), Thyroid carcinoma (THCA), Thymoma (THYM), Uterine Corpus Endometrial Carcinoma (UCEC), Uterine Carcinosarcoma (UCS) and Uveal Melanoma (UVM). C. % euploid (left) and aneuploid (right) tumor samples with p53 deficiency ((–), pink) or proficiency ((+), grey) per cancer type. Hashtags (#) indicate cancer types that have less than 10 euploid samples. The yellow shading depicts pan-cancer aneuploidy prevalence. D. Correlation between % aneuploidy and % p53-deficient samples per cancer type. Grey dashed lines indicate the pan-cancer % aneuploidy (y=86.1%) and pan-cancer % p53-deficient samples (x=38%). Pearson’s correlation test was used. E. Distribution of WGD (dark blue) and non-WGD events (light blue) in tumor samples with p53 proficient ((+), left) or deficient ((–), right). Hashtags (#) indicate cancer types that have less than 10 p53-deficient or -proficient samples. The yellow shading depicts pan-cancer aneuploidy prevalence.

In most cancer types (28 of 31), more than 50% of samples were aneuploid, with a pan-cancer aneuploidy prevalence of 86.1% (Fig. 1B, Suppl. Table 1A). p53 deficiency was significantly associated with aneuploidy (Fisher Exact test: *p* < 0.001, Odds Ratio (OR) = 6.6): the pan-cancer prevalence of p53 deficiency was approximately 38%, with a ∼2-fold enrichment in aneuploid samples compared to euploid ones (40.4% vs 17.8%, respectively) (Fig. 1C, D and Suppl. Table 1A). The enrichment was significant in 10 cancer types (out of 17 eligible for Fisher Exact test (Suppl. Table 4A): BLCA, BRCA, COADREAD, ESCA, LIHC, LUAD, HNSC, PRAD, STAD, UCEC), and these analyses revealed some notable nuances. Although we found p53-deficient euploid samples and p53-proficient aneuploid samples (Fig. 1C), no cancer types with a high prevalence of p53 deficiency (above pan-cancer average of 38%) were predominantly euploid (aneuploidy prevalence above 86.1%) (Fig. 1D), in accordance with the pan-cancer association. However, despite occasionally low sample numbers (Suppl. Table 1A), 7 of 17 cancer types (LGG, PAAD, SARC, SKCM, THCA, THYM, UCS) showed no significant difference between the prevalence of p53 deficiency in euploid and aneuploid samples (Fig. 1C). Several cancer types (e.g. CESC, GBM, KIRC, KIRP, PCPG, SARC, SKCM, TGCT) had an aneuploidy prevalence above 90% but a frequency of p53 deficiency that was below the pan-cancer average (Fig. 1C, D). Similarly to previous reports (*24, 25*), we observe that WGD events in the cancer samples were significantly associated with p53 deficiency (Fisher Exact test: p < 0.001, OR = 3.1) (Fig. 1E and Suppl. Table 3A). Nevertheless, this was also nuanced by a lack of a significant association in ∼70% of cancer types (17 of 24 cancer types eligible for Fisher Exact test, Suppl. Table 4B). In summary, p53 deficiency is significantly associated with aneuploidy at a pan-cancer level, as noted before, but the association was weak or absent in ∼40% of cancer types.

### A high degree of aneuploidy is generally but not necessarily associated with p53 deficiency

While we show that the association of p53 deficiency with aneuploidy is highly cancer-type specific, certain features of aneuploidy might be more widely associated with p53 deficiency. One of these features is the degree of aneuploidy above which p53 inactivation might be required to sustain proliferation (*17, 19, 26*). To examine this, we quantified the degree of aneuploidy by calculating the fraction of genome altered (FGA) in all non-WGD aneuploid tumors (awCNA tumors, carrying arm-level and/or whole chromosome CNAs, see Fig. 2A and Methods). The FGA was agnostic of direction of the alterations unless stated otherwise. For group-level comparisons, average FGA values were used across cancer types and in pan-cancer analyses. Unless specified, WGD tumors were excluded from our analyses to remove the effect of the high aneuploidy characteristic of WGD tumors.

**Figure 2.**
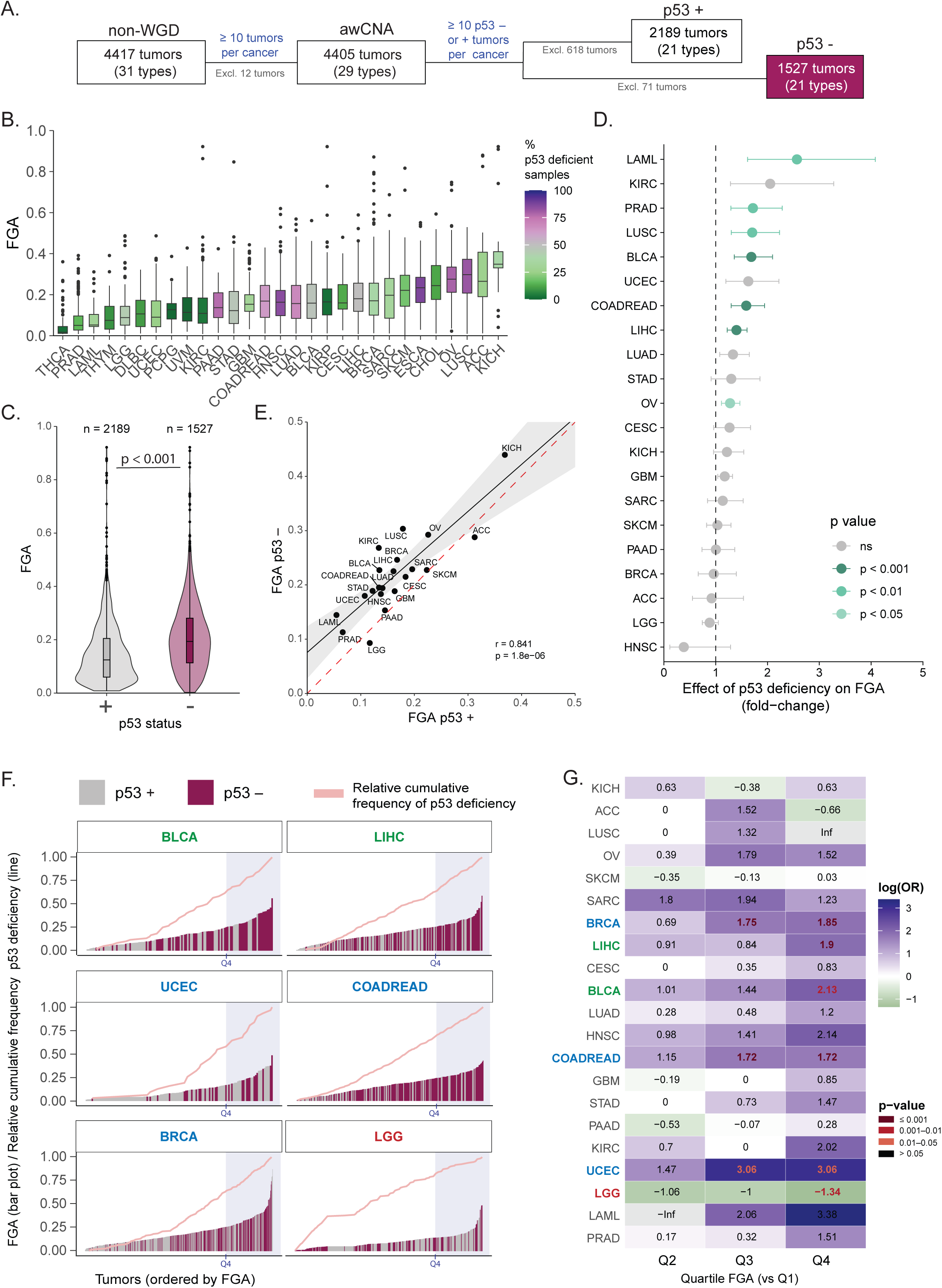
A high degree of aneuploidy is generally but not necessarily associated with p53 deficiency. A. Schematic representation of filtering applied to the non-WGD samples. awCNA corresponds to all possible combinations of whole arm-level and/or whole chromosome CNA events (awCNAs). P53 status: deficient (–) and proficient (+). B. Fraction of genome altered (FGA) for awCNA events per cancer type (ordered by mean FGA). Color filling depicts the % of p53-deficient samples per cancer type. C. Pan-cancer FGA distribution in p53-proficient (grey) and p53-deficient (pink) tumors. Boxplot represents the median and first and third quartiles. Dots represent outliers. “n=” number of tumor samples per group. Statistical analysis was performed using Mann-Whitney-U test. D. Fold-change of p53 deficiency (β_p53_, Gamma GLM) on FGA of tumors, per cancer type=. Colors indicate the significance of the effect: bars are colored by p-value threshold (p<0.05, p<0.01, p<0.001) and shown in grey when the effect is not significant (ns). Error bars represent the standard error of the model coefficient, accounting for overdispersion in the Gamma GLM framework. E. Distribution of FGA in p53-proficient and p53-deficient tumors, per cancer type. Linear regression line (black) and confidence interval (grey shade). The doted red line depicts a 1:1 ratio, corresponding to ratio = 1 as used in D. Pearson’s correlation test was used. F. Distribution of tumor samples in ascending order of FGA for cancer types BLCA, LIHC, UCEC, COADREAD, BRCA and LGG. The color filling corresponds to p53 status *–* proficient (–) (grey) and deficient (+) (pink). Line graphs (light pink) show the relative cumulative frequency of p53-deficient samples. Shaded regions (light blue) correspond to the fourth quartile for each cancer type. Cancer types are colored based on the significantly different distributions of p53 deficiency in relation of the degree of aneuploidy (calculated in B.). G. Log odds ratio of FGA in p53-deficient tumors in quartile 2, 3 and 4 compared to quartile 1. Color coded numbers depict statistical significance calculated with pairwise Mann-Whitney U test (Bonferroni correction method). Highlighted cancer types are colored based on the different distributions of p53 deficiency in relation of the degree of aneuploidy.

Similar to a previous analysis (*4*), aneuploid tumors showed a large variability in FGA within a cancer type and across cancer types (Fig. 2B). Likewise, the prevalence of p53 deficiency varied widely between cancer types with similar average FGA distribution (Fig. 2B). At a pan-cancer level, p53-deficient tumors had a significantly higher average FGA than p53-proficient ones (0.21 and 0.15, respectively; Man-Whitney U test: p < 0.001) (Fig. 2C). However, for the majority of cancer types (66.6% (14/21) there was no significant association of p53 deficiency with FGA when adjusted for age, sex, subtype and stage as confounders (when applicable; see Methods section) (Fig. 2D, Suppl. Table 2B; Suppl. Table 5). Strikingly, for kidney chromophobe (KICH) and adrenocortical carcinoma (ACC), the average FGA could be very high (>0.25) regardless of p53 status (Fig. 2E). We further validated our results by using arm CNA counts (same as the ‘aneuploidy score’ defined by Taylor *et al* (*3*)), which directly correlated with FGA and yielded comparable associations (Suppl. Fig. 3). In general, the association of p53 deficiency and average FGA was independent of chromosome identity for any given cancer type, except for gain of 12p arm enriched in p53-deficient HNSC tumors and gain of 20p enriched in p53-proficient KIRC tumors (Suppl. Fig. 4). In addition, p53 was associated with aneuploidy in estrogen receptor-positive breast cancer but not in estrogen-receptor-negative breast cancer (1.52-fold higher FGA; Gamma GLM: β_p53*subtype_= 0.489, *p* = 0.018) (Suppl. Fig. 5). For all other cancer types, p53 was associated with higher FGA but this was not significantly impacted by cancer subtype (Suppl. Fig. 5). WGD samples had a similar FGA distribution in the majority of cancers independently of p53 status (Suppl. Fig. 1C,D, Suppl. Table 4F). Analysis of oncogenic mutations in p53-proficient tumors showed enrichement of only four genes (PBRM1, VHL, CIC and SPOP), all of which had low pan-cancer prevalence; even those of the p53 pathway (CDKN1A, ATM and MDM2/4) yielded none that was consistently enriched across cancer types (Suppl. Fig. 6).

### Tumors with the highest degree of aneuploidy are not significantly enriched for p53 deficiency

Although for two thirds of cancer types the average FGA was not significantly different in p53-proficient and deficient samples, we next wondered whether within a cancer type p53 deficiency might be more prevalent among the samples with the highest FGA. To examine this, we ranked all samples within a cancer type from low to high FGA and assessed p53 status in the four quartiles (Fig. 2F,G and Suppl. Fig. 7). We found that, in the majority of cancer types (15 of 21), the samples with the highest FGA (top quartile) were not significantly enriched for p53 deficiency when compared to those with the lowest FGA (bottom quartile) (Fig. 2G, Suppl. Fig. 7 and Suppl. Table 4C). This was independent of the percentage of p53-deficient samples per FGA quartile (Suppl. Fig. 7B). From the seven cancer types with a significantly higher average FGA in p53-deficient samples (Fig. 2D), only two showed an enrichment specific to the top FGA quartile (BLCA and LIHC). Three cancers showed a significant enrichment in the top two quartiles (BRCA, COADREAD and UCEC). On the contrary, LGG was enriched for p53-deficient samples with low FGA (Fig. 2F,G).

These analyses show that tumors with high aneuploidy yet intact p53 activity are common, and that p53 inactivation does not invariably lead to highly aneuploid tumors. p53 inactivation is therefore neither sufficient nor necessary for tumors to accumulate a high degree of aneuploidy.

### p53-deficient cancers are enriched for chromosome losses

Having found no universal association between p53 deficiency and the degree of aneuploidy, we next investigated whether there might be an association with the type (arm-level or whole chromosome) or direction (gains or losses) of aneuploidy. Isolating samples with only arm-level CNAs (aCNAs) or whole-chromosome CNAs (wCNAs) did not show a significant association of type-specific FGA with p53 status, beyond what could be explained by generally elevated FGA (Suppl. Fig. 8).

Previously, a correlation was found between p53 loss and a below-diploid genome content (*18*). To examine a possible association of p53 status with the direction of aneuploidy, we categorized the tumor samples with both arm-level and whole chromosome CNAs (bCNAs, 2530 samples) as having a gain or loss when at least one arm or whole chromosome was amplified (2015 samples) or deleted (2125 samples), respectively (Fig. 3A). We then calculated the fraction of the genome affected by losses (FGA-L) or gains (FGA-G) in each tumor sample. Note that this was agnostic of total FGA (excluding WGD), which could on average be negative or positive and therefore does not reflect whether tumors were below or above diploidy. We observed no difference in the fraction of the genome gained between p53-deficient or -proficient cancers, neither at the pan-cancer level nor in most cancer types (Fig. 3B and Suppl. Fig. 9A,B, Suppl. Table 4D). Notably, however, most p53-deficient cancers had a higher fraction of the genome lost than did the p53-proficient cancers (significant for 6 out of 17 cancer types: Fig. 3C; Suppl. Fig. 9A,B and Suppl. Table 4E). No significant difference was observed in cancers that had undergone WGD (Suppl. Fig. 9F,G; Suppl. Table 4G and H). These analyses therefore indicate that p53 inactivation is specifically associated with an increased incidence of chromosome losses but not gains. Of note and consistent with a previous report (*18*), p53 deficient cancers had a consistently higher fraction of the genome lost than gained across cancer types (significant for OV and UCEC: Fig. 3D, and Suppl. Fig. 9B-E, 10, 11), resulting in a below-diploid genome content.

**Figure 3.**
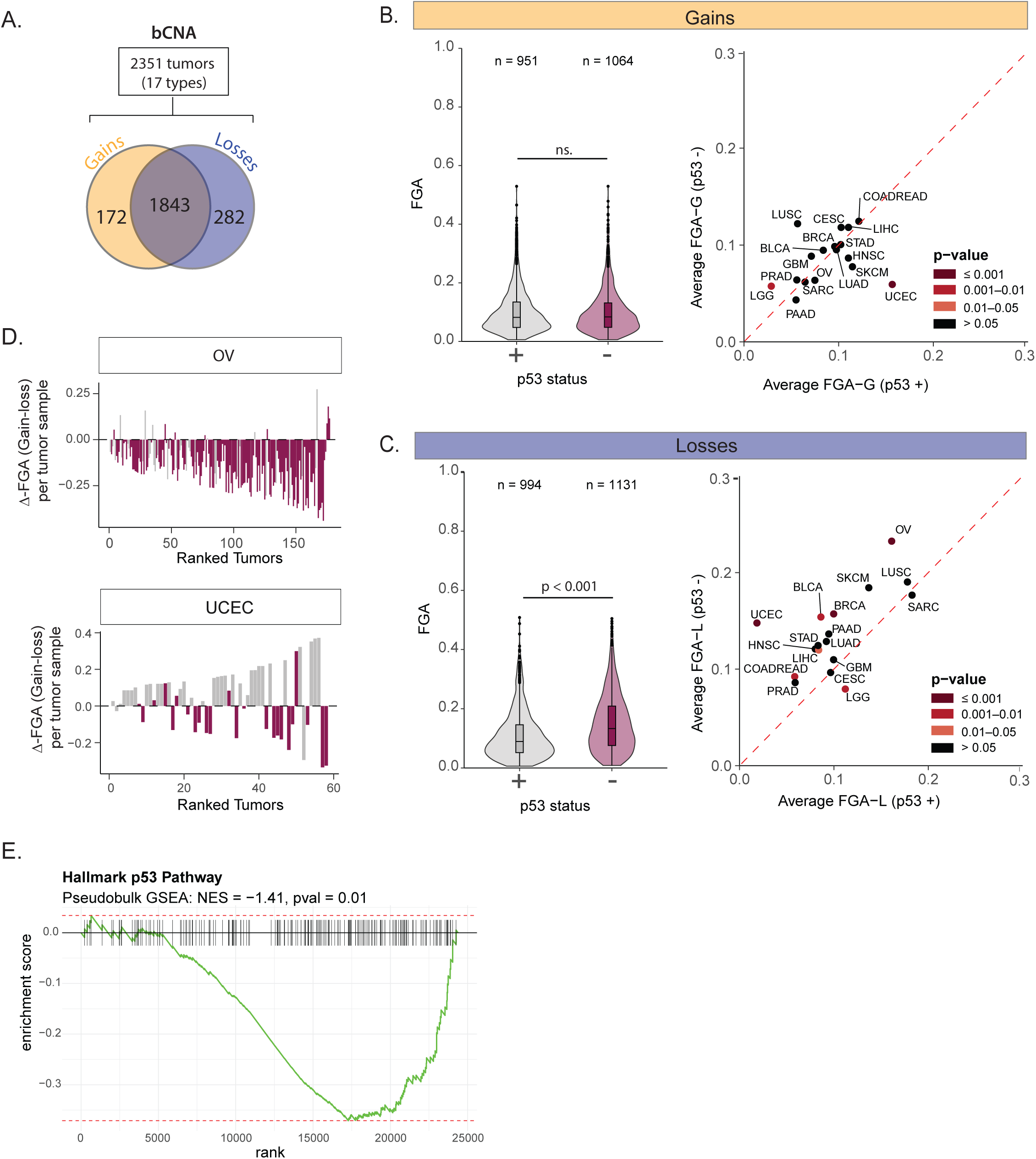
Chromosome losses are more prevalent in p53-deficient cancers. A. Categorization of gain (yellow) and loss (blue) copy number events in bCNA samples. Copy number gain events include samples with only gain events (172 samples) or gains +losses (1843). Copy number loss events included for samples with only loss events (282 samples) or gains +losses (1843). B. Distribution of FGA attributed to gains (FGA-G). Left: Pan-cancer FGA distribution of bCNAs in p53 deficiency (grey) and proficiency (pink). Right: Distribution of FGA attributed to gains in p53-proficient and p53-deficient tumors, per cancer type. Doted red line depicts a 1:1 ratio. Cancer types are colored based on the significant distributions of p53 deficiency in relation of the degree of FGA-G; pairwise Mann-Whitney U test (Bonferroni correction method). C. Same as in C. for FGA attributed to losses (FGA-L). D. Delta of FGA (FGA-G minus FGA-L) per tumor sample of Ovarian (OV) and Uterine and Endometrial (UCEC) cancer types. Samples are distributed in ascending order of total FGA. The color filling corresponds to p53 status: proficient (+) (grey) and deficient (–) (pink). Positive values correspond to the higher burden of FGA-G and negative values to a higher burden of FGA-L in each tumor sample. E. Pre-ranked GSEA plot demonstrating the suppression of the Hallmark p53 Pathway in response to an increasing chromosomal loss fraction. The analysis was performed on pseudo-bulk expression data across the tumor cohort. To strictly control for known confounding variables, the gene ranking metric was derived from the coefficients of a multivariable linear regression model, which evaluated gene expression against the continuous loss fraction while adjusting for total malignant aneuploidy score, tumor purity, and cancer lineage. The negative Normalized Enrichment Score (NES = **-1.41**, nominal *p* = **0.01**) indicates that as the loss fraction increases, the global p53 transcriptional program is significantly downregulated relative to the rest of the genome.

To further confirm these findings using an independent dataset, we analyzed matched aneuploidy profiles and gene expression from a recent resource of single cell RNAseq data from ∼288,000 malignant cells, across 304 tumors and 15 cancer types (*27, 28*). At the tumor level, we found a significant association between the prevalence of losses (as a fraction of all CNAs) and the downregulation of the p53 pathway (Fig. 3E).

### p53 inactivation selectively enriches for chromosome losses in non-transformed cell lines

Thus far, our analysis showed an association between p53 deficiency and higher occurrence of chromosome losses. To examine whether p53 deficiency causes selective enrichment of losses, we examined the karyotype of four sets of p53-proficient (*TP53* WT) and -deficient (*TP53* KO) cell lines: two non-transformed lines (RPE-1 (retinal pigment) and MCF10A (breast)) and two cancer lines (A549 (lung) and HCT116 (colorectal)). Single-cell karyotype sequencing (scKaryoSeq, (*29*)) showed that all p53-proficient lines displayed clonal aneuploidy patterns characterized mostly by chromosome gains (Fig.4A,B and Suppl. Fig. 12A-F). We observed a significant increase in FGA in *TP53* KO cells compared to *TP53* WT in both non-transformed cell lines, but not in the cancer cell lines (Suppl. Fig.12G). Strikingly, while none of the p53-proficient vs -deficient cell lines had a significant difference in the fraction of the genome gained, the two non-transformed lines had a significant, selective increase in the fraction of the genome lost after *TP53* deletion (Fig. 4C,D). Across chromosome arms, the proportion of cells harboring loss events was two and four-fold higher in *TP53* deleted RPE-1 and MCF10A cells, respectively, while gains were restricted to the clonal alterations (Fig. 4E and Suppl. Fig. 12H).

**Figure 4.**
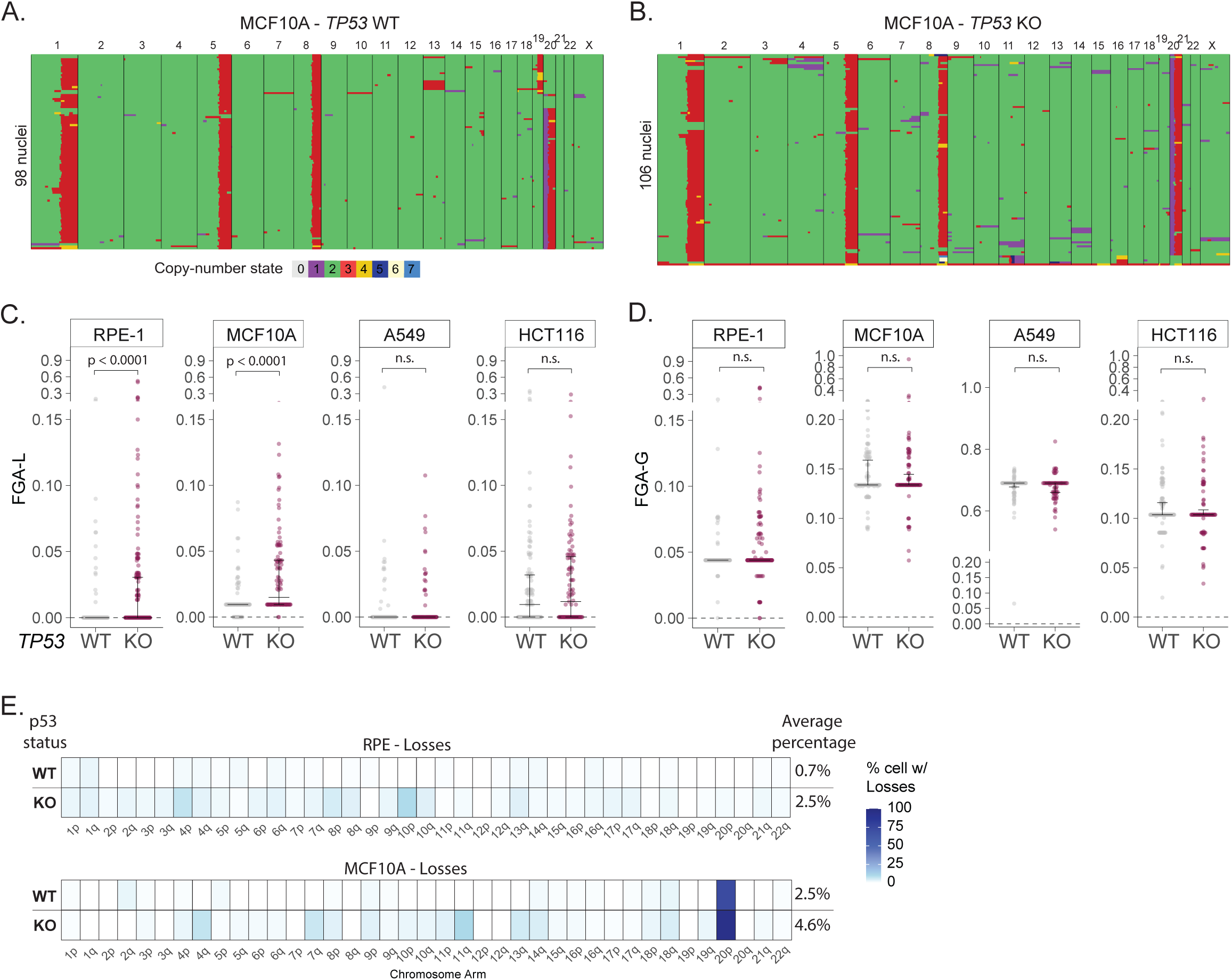
p53 inactivation selectively enriches for chromosome losses in non-transformed cell lines. A-B. Representative single cell karyotype sequencing plots for MCF10A cell line. Conditions: TP53 WT (A) and TP53 KO (B). Color code depicts copy number states. Left: Number of sequenced nuclei. C. Distribution of FGA attributed to losses (FGA-L) across cell lines ((RPE, MCF10A, A549 and HCT116) with TP53 WT (grey) or KO (pink). Data corresponds to 2 to 3 replicates for RPE-1 cells and one replicate for the other lines. Kruskal-Wallis test with pairwise Mann-Whitney U tests and Bonferroni correction for multiple comparisons was used. D. Same as in C. for FGA attributed to gains (FGA-G). E. Average frequency of loss events per chromosome arm in RPE-1(top) and MCF10A (bottom) cells with TP53 WT or KO. Right: average percentage of cells with loss events per condition.

### Induced chromosome losses activate the p53 response

The results thus far suggest that p53 is a selective barrier to survival and proliferation of cells that have lost chromosomes. To test this more directly, we induced chromosome segregation errors in RPE-1 cells and assessed the p53 response to gain or loss events and the surviving karyotypes after long-term culture (Fig 5A). Single cell RNA sequencing of p53-proficient cells following induced segregation errors enabled inference of chromosomal copy number states and p53 pathway activation (Fig 5A-1; See Methods section). Cells that had exclusively lost whole chromosomes (one or more events per cell) had a significantly higher p53 pathway activation than cells with exclusively chromosomal gains (Fig. 5B). This was consistent across cells with different degrees of gains or losses, even very low (FGA 0.01-0.05) (Fig. 5C). Moreover, single loss events but not single gain events were sufficient to consistently induce a p53 response (Suppl. Fig 13).

**Figure 5.**
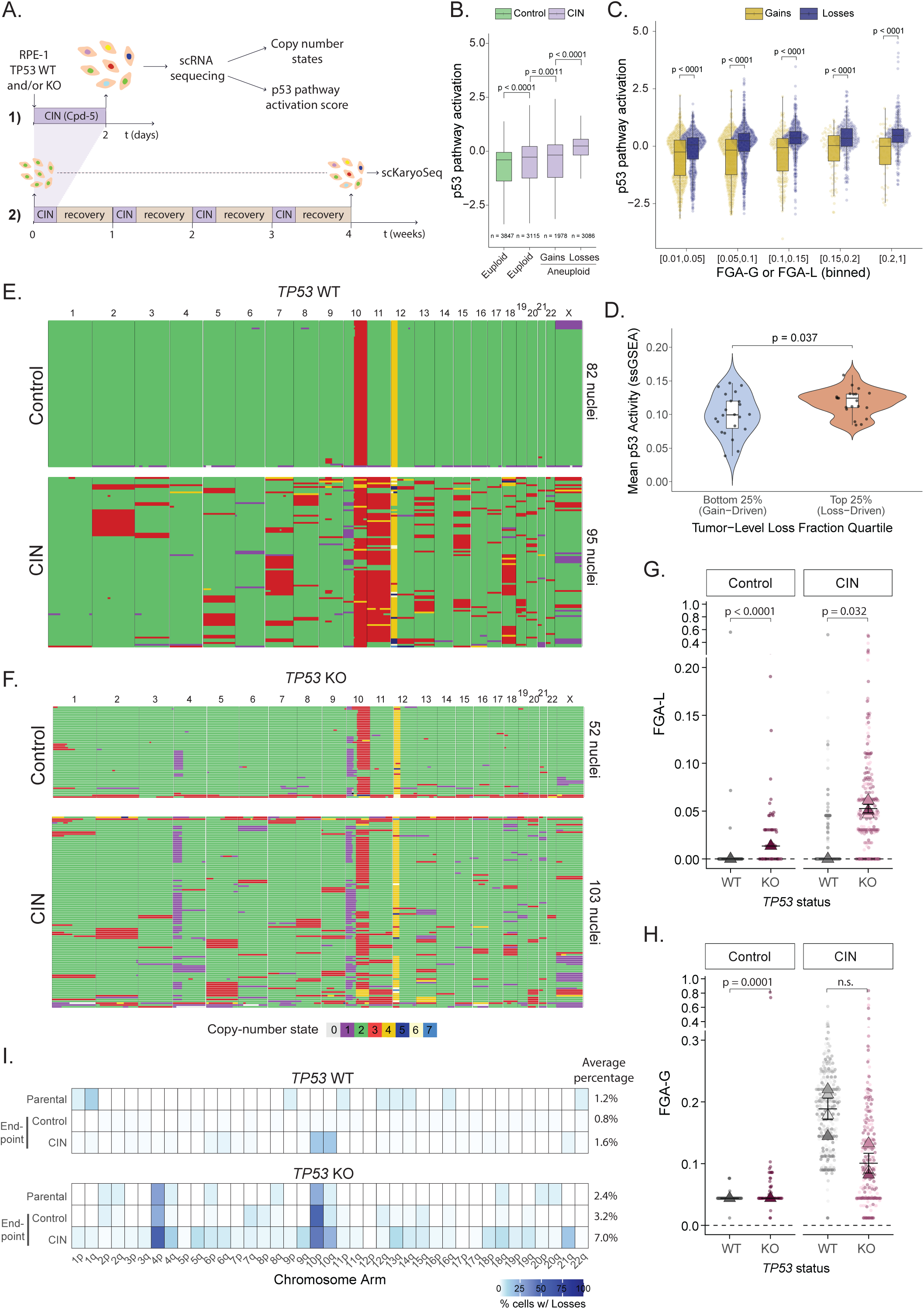
p53 inactivation bias for loss events long-term in non-transformed RPE-1 cell line. A. Schematic representation of experimental set-up to assess short- and long-term p53 response to loss events. 1) Short-term CIN induction (Cpd-5, (*51*)) for 2 days followed by collection of cells for single cell RNA sequencing (scRNAseq), in *TP53* WT RPE-1 cells. Copy number states and p53 pathway activation scores were inferred from scRNAseq. 2) 4 cycles of intermittent treatment with Cpd-5 for 2 days followed by 5-day recovery period, in continuous cultured *TP53* WT and KO RPE-1 cells. Cells were collected for scKaryoSeq at the start- and endpoint (Parental and 4 weeks, respectively). B. p53 pathway activation score for euploid (FGA = 0) control (green) and euploid, whole-chromosome gains only (at least one event per cell) and whole-chromosome losses only (at least one event per cell) CIN-induced cells (purple). Box plot represents median and first and fourth quartile. n = the number of cells per condition. Student’s t test was used. C. p53 pathway activation score for cells with only whole-chromosome gains (at least one event per cell; yellow) or only whole-chromosome losses (at least one event per cell; blue), binned per increasing degree of FGA. Each dot represents one single cell. Box plot represents median and first and fourth quartile. Student’s t test was used. D. Violin plot comparing the mean p53 pathway activity (ssGSEA) between extreme tumor-level loss fraction quartiles. The analysis was restricted to a subset of tumors exhibiting high karyotypic heterogeneity (AHS ≥ upper tertile). Tumors in the bottom 25% (Gain-Driven, blue) are compared against those in the top 25% (Loss-Driven, orange). Boxplots represent the interquartile range with the median indicated by the solid central line; individual dots represent the mean ssGSEA score of malignant cells for a single tumor sample. Statistical significance was determined using an unpaired, two-sided Wilcoxon rank-sum test (*p* = **0.037**). E,F. Representative single cell karyotype sequencing plots for RPE-1 cell line. Conditions: *TP53* WT (E) and *TP53* KO (F); control (top) and CIN (bottom). Color code depicts copy number states. Right: Number of sequenced nuclei. G. Distribution of FGA attributed to losses (FGA-L) in RPE with *TP53* WT (grey) or KO (pink), for control and CIN conditions at endpoint (4 weeks). Each dot represents an individual cell, colored by replicate, when applicable. *TP53* WT (shades of grey) and *TP53* KO (shades of pink). Triangles indicate the median value per replicate. Horizontal bars represent the mean of replicate medians ± standard error of the mean (SEM). Control has only one replicate; the bar represents the median of all cells ± SEM. Statistical comparisons between WT and KO within each condition were performed using a two-sided Mann-Whitney U test. For control condition, all individual cells were used as observations. For CIN condition, replicate medians were used with each replicate representing an independent experiment. H. Same as in G. for FGA attributed to gains (FGA-G). I. Average frequency of losses per chromosome arm in RPE-1 cells with *TP53* WT or KO, for parental and endpoint conditions (control and CIN). Right: Average percentage of cells with loss events per condition.

We next again turned to the single cell RNA seq resource from 304 tumors (*27, 28*). In that dataset, high karyotype heterogeneity was found to be associated with signatures of reduced proliferation, acute stress, and p53 pathway activation (*28*). We therefore hypothesized that these signatures might be driven by chromosomal losses. Strikingly, the samples with the highest loss fraction (out of total CNAs; referred to as “loss driven”) in that subset of karyotypically heterogenous tumors had significantly higher p53 pathway activation than those with the lowest loss fraction (“gain driven”) (Fig. 5D), verifying in tumor samples what we have observed in the CIN-induced RPE-1 cells.

We next exposed *TP53* WT and *TP53* KO RPE-1 cells to a four-week regimen of intermittent chromosomal instability followed by recovery (Fig.5A-2). Both *TP53* WT and KO cells showed an initial proliferation decline after one week (∼50% and ∼70%, respectively), with full recovery of the *TP53* KO cells by four weeks (Suppl. Fig. 14F,G). scKaryo-seq showed that while both p53-proficient and -deficient cells had higher FGA after the CIN regimen compared to control conditions (Fig. 5D,E; Suppl Fig. 14A,B), p53 deficiency led to a strong and selective enrichment of chromosome losses, reflected by a significant increase in FGA-L (Fig. 5F). Additionally, a substantially larger fraction of CIN-exposed *TP53* KO vs WT cells harbored at least one chromosome arm loss event (7.0% vs 1.6%) (Fig 5H). Of note, *TP53* WT cells showed high aneuploidy levels, almost exclusively attributed to chromosomal gains (Fig. 5G and Suppl. Fig. 14E,F).

## Discussion and conclusions

p53 inactivation and aneuploidy play central roles in tumorigenesis and prior studies have linked *TP53* mutations to cancer aneuploidy (*3, 13, 16*). Our analysis builds on this by defining various types of aneuploidy and analyzing these in the light of p53 status, for 31 cancer types in TCGA. We find that p53 deficiency is associated with the degree of aneuploidy only in a limited number of cancer-types, and that it is more universally associated with chromosome losses. *In vitro* examination of this association showed selective activation of the p53 pathway to chromosome loss events, and the selective survival of cells with chromosome losses only when p53 was inactivated.

Mildly aneuploid cells in culture can evade p53 surveillance (*19, 22, 26, 30*) while highly aneuploid cells often require p53 inactivation to proliferate (*31*). The relatively low degree of aneuploidy in many tumors may thus fall below the threshold required to activate p53. However, we observed highly aneuploid tumors proficient for p53 across multiple cancer types, challenging the presence of a universal p53-mediated stress response to aneuploidy. Instead, we find that chromosome losses, but not gains, strongly correlate with p53 deficiency across cancer types, irrespective of the degree of aneuploidy in the tumor samples. This agrees with a study reporting enrichment of p53 pathway deficiency in cancers with a below-diploid genomic content (*18*). Moreover, in a mouse model of pancreatic cancer progression, p53 loss is followed by preferential and clonally homogeneous accumulation of chromosome losses (*32*). Selective surveillance for chromosome losses may thus contribute to the role of p53 in tumor suppression. In line with this, we found that all levels of chromosome loss triggered a p53 response, whereas the response to chromosome gains was more variable and scaled with the fraction of the genome affected. This asymmetry suggests that gains and losses impose distinct cellular stresses that differentially act on p53. A possible explanation for stronger p53 activation upon chromosome loss vs gains is haplo-insufficiency of genes involved in vital cellular pathways (*33*). The dosage imbalance caused by losses may be more pronounced than those caused by equivalent gains. One such pathway is ribosome biogenesis, which was shown to be affected by various monosomies (*18*) and which can cause p53 activation (*34*). Uncovering which stresses in response to which chromosome losses cause p53 activation and whether these are universal for tissues will be an important avenue for future research.

Our observation that p53 deficiency is not required for the emergence of high aneuploidy raises the question of how cells adapt to aneuploidy when the p53 response is intact. Besides direct alterations to the p53 pathway, tumors might develop mechanisms of aneuploidy tolerance that act independently of p53, such as p38 inhibition (*35*), BCL-9L inactivation (*36*), or overexpression of cyclin D1/2 (*37, 38*). Nonetheless, the low pan-cancer prevalence reported for these alterations suggest that important other unknown aneuploidy adaptation mechanisms are likely to exist (*36, 37, 39*). Additional adaptive responses may include increased activity of DNA repair pathways (such as DNA damage response and MiDAS), the RAF/MEK/ERK signaling cascade, and RNA or protein degradation mechanisms, all of which have been reported in p53-proficient aneuploid cells (*26, 40, 41*). The cancer type-specific association between p53 deficiency and aneuploidy suggests a strong dependency on tissue context, for example in the types or degrees of stress responses to aneuploidy. In line with this, cell type-specific p53 dynamics have been shown to elicit distinct responses to DNA damage (*42-46*), which may underlie variable susceptibility to broader aneuploidy-induced stresses (*47*). Factors such as differential tissue susceptibility or sensitivity to nuclear deformations (*48*) or tissue-specific metabolic profiles (*49*) might also influence the degree of p53 activation in response to aneuploidy. Furthermore, distinct *TP53* mutations may exert differential effects depending on the cancer type (*50*), underscoring the need for future studies to clarify their role in the emergence of aneuploidy. Understanding the tissue context of p53 responses to aneuploidy and uncovering p53-independent mechanisms for aneuploidy tolerance will require in-depth studies using appropriate models of diverse healthy and pre-cancer human tissues.

## Acknowledgments

We thank members of the Kops lab for helpful discussions. OpenAI’s ChatGPT-4o was used to assist in the development and debugging of R scripts employed in the analysis presented in this manuscript. The Kops lab is part of Oncode Institute, which is partly funded by the Dutch Cancer Society (KWF Kankerbestrijding). This work was supported by the Dutch Cancer Society (project 12728) and by the European Research Council (ERC-SyG 855158). Work in the Ben-David lab is funded by the Israel Cancer Research Fund Project Award, the Israel Science Foundation project grant (1805/21), the U.S.-Israel Binational Science Foundation (BSF) project grant (2019228), the European Research Council Starting Grant (945674), and the Schmidt Science Polymaths Award.

## Author contribution

Conceptualization: J.F.M., G.J.P.L.K. Formal analysis: J.F.M., M.D.L., G.W.D., M.S. Investigation: J.F.M., T.W. Methodology: J.F.M., M.D.L., G.W.D., M.S. Resources: T.D., U.B.D. Writing – original draft: J.F.M., G.J.P.L.K. Writing – review & editing: all authors.

## Data, code, and materials availability

All code can be access in the Github - https://github.com/jrfmarques/TCGA_p53_aneuploidy. All data used in this manuscript is available in ENA repository - PRJEB114337.

## Methods

### 1. Dataset

We analyzed 8539 TCGA tumor samples across 31 cancer types with available data for copy-number alteration (CNAs), ploidy, and p53 status. See Suppl. Table 1 and 2 for total sample numbers and CNA counts and FGA calculations. We did not include human papillomaviruses (HPV)-positive HNSC tumors because HPV directly promotes p53 degradation (*52*). PAAD tumors previously identified as misclassified in TCGA were excluded from the analysis (*53*).

#### 1.1. CNA calling

Somatic DNA copy number profiles were obtained from a published dataset filtered for whole-arm and whole chromosome CNAs (*4, 54*). DNA copy number were determined from Affymetrix 6.0 SNP arrays using GISTIC 2.0 from TCGA (version 2016/01/28) and recalculated based on tumor purity estimates using ABSOLUTE (version 1) or pathology reports (*55-57*), and assuming an average ploidy of 3. Briefly, arm-level CNAs were called when a change in DNA copy number spanned more than 98% of the chromosome arm. In addition, we applied a threshold of +/- 0.3 to the copy number value for filtering true CNAs. Whole chromosome CNAs were called when the copy number value of both the p and q arms were equal or considered 0 if otherwise. This threshold can be interpreted as 30% of the tumor cells, on average, in the population carrying said CNA. Acrocentric chromosomes (chr. 13, 14, 15, 21 and 22) were considered as whole chromosomes. For CNA counts, alterations were considered +1 when amplified, -1 when deleted, and 0 when unaltered. For the calculation of fraction of genome altered (FGA), CNAs were considered to be the fraction of the genome for each arm based on its length in base pairs (extracted from reference genome hg38), positive when amplified and negative when deleted.

#### 1.2. WGD filtering

Whole Genome Doubling (WGD) data was extracted from ABSOLUTE version 1 (publicly avalaible) (*58*), and matched to the current dataset. WGD events >0 (either 1 or 2) where grouped as positive WGD samples. Of note, this data was not used to derive the copy-number states used in this work.

#### 1.3. Mutations calling

Tumor samples were considered as having *TP53* mutations when any number of nonsense, splice site, frame shift, or missense mutations were called. Missense mutations were considered only if Polyphen2-HVAR score > 0.2, which predicts the impact of amino acid substitution on protein structure and function. When these were absent, samples were classified as wild type for *TP53. TP53*-loss phenocopy callings were obtained from (*23*). These callings were used to re-call *TP53* wild type samples as p53-deficient samples. This classification includes alterations to the p53 pathway, namely amplifications of MDM2/4 and PPM1D, deletion of *TP53* and copy number alteration of MDM2/4, PPM1D and *TP53*. A total of 5035 tumors were present in our dataset and from these, 8% were classified as *TP53*-loss phenocopy. Mutations in significant driver genes in cancer were derived from (*59*). Oncogenes and tumor suppressor genes with a TUSON *q*-value smaller than 0.05 were selected.

#### 1.4. Data filtering

From the data set, 1144 samples were euploid and 7395 samples were aneuploid, of which 40% (2978) were designated as having undergone a whole genome doubling (WGD). The three categories (euploid, aneuploid +/- WGD) were further subdivided into samples proficient or deficient for p53 (see “Mutations calling” section). Cancer types with fewer than 10 aneuploid tumor samples were excluded from the analysis. For comparisons of p53 status in aneuploid samples, we excluded cancer types with fewer than 10 p53-deficient samples. The category of all aneuploid samples (awCNA) were further subdivided into: aCNAs – only whole arm-level CNAs (if arm>0; whole=0); wCNAs – only whole chromosome CNAs (if arm=0; whole>0); bCNAs – both whole chromosome and whole arm-level CNAs (if arm>0; whole>0). Of note: the latter category can include any combination of gain or loss events as long as at least one of each CNV type (whole chromosome or whole arm) events are present. Samples were classified as having gains if they included at least one whole-chromosome or arm-level CNA with a positive FGA/count, and as having losses if they included at least one event with a negative FGA/count.

### 2. scRNA sequencing tumor data resource

Single-cell RNA sequencing (scRNA-seq) data were compiled from 304 tumor samples across 36 independent studies, spanning 15 distinct cancer types. This integrated dataset was sourced from the Curated Cancer Cell Atlas (3CA), which provided both the normalized gene expression profiles and the associated cell-level annotations (*27*). To establish the karyotypic landscapes of the single cells, gene-level copy number estimates were computationally inferred from the transcriptomic data using the InferCNA algorithm. These gene-level matrices were subsequently processed using the ASCETS algorithm to generate robust chromosome arm-level copy number calls. This framework enabled the calculation of absolute Aneuploidy Scores (AS) — defined as the total number of gained or lost chromosome arms — and the extraction of specific structural loss and gain fractions for every individual malignant cell in the cohort (*28*).

#### 2.1. Pseudo-bulk level analysis

To investigate the continuous relationship between chromosomal loss burdens and global pathway activity, pseudo-bulk differential gene expression (DGE) analysis was performed. For each tumor sample, a Loss Fraction was calculated as the ratio of the mean malignant chromosome loss score to the overall malignant mean aneuploidy score. This analysis was restricted to a subset of 255 tumor samples that retained a sufficient number of malignant cells (>=50) to reliably calculate the tumor-level loss fraction.

##### 2.1.1. Gene-Level Multivariable Regression

We calculated the exact Loss Fraction (Malignant Loss Score / Total Malignant AS) for each tumor as a continuous variable (from 0.0 to 1.0). We then fitted a multivariable linear regression model for *every single gene in the transcriptome*. In this model, gene expression is the dependent variable, and the continuous Loss Fraction is the primary independent variable. We also strictly controlled for total malignant AS, tumor purity, and cancer type as covariates.

##### 2.1.2. Pathway -Level Gene Set Enrichment Analysis (GSEA)

We extracted the Loss Fraction regression coefficient for every gene. This coefficient represents the independent ’degree’ to which a specific gene’s expression shifts as a tumor becomes more loss-driven. We used these coefficients to rank the entire genome, and ran a standard pre-ranked GSEA on this ranked list.

#### 2.2. Single-cell level analysis

To evaluate absolute pathway activation at the tumor-level, single-sample GSEA (ssGSEA) enrichment scores for specific ’Hallmark’ pathways (e.g., p53 pathway) were computed per cell and averaged across all malignant cells within each sample to derive a Mean p53 Activity score. To ensure robust estimations and complete data integration, the cohort was restricted to samples that possessed matched single-cell loss and gain scores, valid p53 activity profiles, and a sufficient number of malignant cells (≥50) to accurately calculate the tumor-level loss fraction.

##### 2.2.1. Subset based on intra-tumor heterogeneity of tumor samples

a subset of highly heterogenous tumors (114 tumors) were used as defined in Wolf-Dankovich *et al* (*28*). Briefly, pairwise cell-to-cell similarity scores were computed within each sample based on gene-level copy number profiles of malignant cells. For each tumor sample, the mean of all pairwise correlations were computed to define a Karyotypic Heterogeneity Score. This metric reflects the overall similarity in copy number profiles among malignant cells.

##### 2.2.2. Data stratification

data was stratified into discrete quartiles based on their loss fraction. Tumors in the bottom 25% quartile were defined as "Gain-Driven," while those in the top 25% quartile were defined as "Loss-Driven." Statistical differences in mean ssGSEA scores between these extreme quartiles were evaluated using an unpaired, two-sided Wilcoxon rank-sum test. Data distributions were visualized as violin plots overlaid with boxplots and jittered individual sample points using the ggplot2 and ggpubr R packages.

#### 3. Cell culture and generation of cell lines

RPE1-hTERT (ATCC-CRL-4000), A549 (ATCC-CCL-185) and HCT116 (ATCC-CCL-247) cells lines were cultured in DMEM/F12 with GlutaMAX supplement (Gibco), RPMI (Gibco) and McCoy (Gibco), respectively, and supplemented with 10% fetal bovine serum (FBS, Sigma-Aldrich) and 1% penicillin/streptomycin (Sigma-Aldrich). TrypLE (Gibco) incubation was used for cell splitting. To obtain *TP53* KO, cells were transduced with a lentivirus containing the *TP53* KO gRNA CGGTTCATGCCGCCCATGC, in a lentiCRISPRv2 plasmid (<u>Addgene: lentiCRISPR v2</u>). Transduced cells were selected with 10 μg ml–1 puromycin (Sigma-Aldrich) for 5 days and *TP53* KO RPE-1 cells were further selected with 10 μM Nutlin 3a (Selleck) for 10 days to validate functional loss of p53. MCF10A were cultured in DMEM/F12 supplemented with 5% Horse serum, 20ng/ul final concentration of EGF 0.5mg/ml final concentration of Hydrcortisone, 100ng/ml final concentration of Cholera toxin, 10ug/ml final concentration of Insulin, 5ug/ml final concentration Transferin and 5ml Pen/strep. Cells were transduced with 2sgRNAs against *TP53* in a LentiCRISPRV2 neomycin resistant (<u>Addgene: lentiCRISPRv2 neo</u>): sg1: CGCTATCTGAGCAGCGCTCATGG; sg2: CCCCGGACGATATTGAACAA. Control MCF10A *TP53* WT cells were transduced with the following empty vector – PLKO1 blast resistant, hPGK promoter (<u>Addgene: pLKO.1-blast</u>). Validation was done on the population via qPCR for p21 and p53.

#### 3.1. Induction and long-term treatment of RPE-1 cells

We created a heterogeneous population of aneuploid cells by applying the following approach: 4 rounds of MPS1 inhibition (Compound 5, Cpd-5 62.5 μM; (*51*)) followed by washout and recovery for 5 days; Cells were plated in 24-well plates (Sigma-Aldrich), and passaged at a ratio of 1:6. Cells were collected before start (parental), at 1-week timepoint for EdU assay and at 4-week timepoint (endpoint) for EdU assay and scKaryoSeq.

#### 3.2. EdU proliferation assay and flow cytometry

RPE-1 cells were plated 1 day before treatment with 10 μM EdU (Thermofisher) for 24h, and fixed using 70% ice-cold ethanol for 1h. Ethanol was removed by one wash with PBS, and cells were incubated for 20 min with the Click-iT reaction cocktail (Click-iT EdU proliferation assay). The reaction cocktail was washed away and replaced with a PBS/DAPI mix. Single cell suspensions were analyzed on a BD LSR Fortessa X-20 (BD Biosciences). Quantification and plotting were performed in FlowJo v10.

### 4. Single-cell karyotype sequencing

*For RPE-1, A549 and HCT116:* cells were trypsinized and stored at −20°C for further processing. Single G1 nuclei were isolated with Nuclei Suspension Buffer (NSB) 100 mM Tris-HCl pH 7.5, 154 mM NaCl, 1 mM CaCl2, 0.5 mM MgCl2, 0.2% BSA, 0.1% NP40, 1 ug/mL Hoechst 34580 and sorted using a BD FACSAria Fusion flow cytometer in Hard-Shell 384-well PCR plates (Bio-Rad). Plates were stored at −20°C. NlaIII-based library preparation was performed as described previously, with several modifications (46). Cell lysis was performed for 2 h at 55 °C with 8 mg ml–1 Proteinase K (Fisher Scientific) in 1× CutSmart (New England Biolabs) and heat inactivation at 80°C for 10 min. NLAIII incubation for 2 hours at 37°C followed by heat inactivation at 65C for 20 min, Adaptors were ligated with 50 nl of 100 nM barcoded, double-stranded NLAIII adaptors and 150 nl of 10 U T4 DNA ligase (New England Biolabs) in 1× T4 DNA ligase buffer (New England Biolabs), supplemented with 3 mM ATP (Invitrogen) at 16°C overnight. Samples were sequenced on an Illumina NextSeq500 or Nextseq2000 sequencer at 1× 75 or 1× 100 single end base pairs (bp), respectively.

*For MCF10A*: cells were sorted on a BD Influx cell sorter with a 100 uM nozzle size. Samples were incubated with DAPI (1:1000) 15 minutes prior to sorting and the sort was based on FCS/SSC (P1 gate) with daughter P2 gate DAPI negative. Cells were sorted into 384-well plates with 5 μL of mineral oil (Sigma-Aldrich). 500 nL of lysis mix (0.0005 u QIAGEN Protease in NEB Buffer 4) was added to each well and lysis was performed at 55°C overnight followed by heat inactivation for 20 min at 75°C and for 5 min at 80°C. 500nl of Restriction Enzyme mix (0.5 u NlaIII in NEB Cutsmart buffer) was added to each well and restriction was performed for 3 h at 37°C followed by heat inactivation for 20 min at 65°C.

100 nL of 1 uM barcoded double stranded NlaIII adaptor was added to each well. 1100 μL of Ligation mix (200 u T4 DNA Ligase in 1x T4 DNA Ligase buffer supplemented with 3 mM ATP) was added to each well and ligation was performed overnight at 16°C. After ligation, single cells were pooled and library preparation was performed as described in Muraro *et al* (*60*). Libraries were paired-end sequenced on the Illumina nextseq 2000 platform according to the manufacturer’s protocol. Coverage: 4% (16 million reads per plate) USEQ NextSeq2000 P2 2x150bp.

#### 4.1. Single-cell karyotype sequencing analysis

Sequences were mapped to the homo sapiens genome (BSgenome H.Sapiens UCSC hg38) using Burrows-Wheeler Aligner (BWA). Aneufinder (v.1.34.0) was performed to obtain copy number states using the e.divisive method. We applied an extra filtering on the Aneufinder output for quality control (https://github.com/TWvR/AneufinderFileFilter) using the following parameters: Minimum total reads – 25000; Maximum spikiness: 0.25; Minimum level Bhattacharyya: 0.7. Copy number variations of whole and partial chromosomes were determined using R scripts developed in the lab, R (v4.4.2) and R-Studio (v 2024.09.1).

#### 4.2. FGA calculation

From absolute CNA counts obtained from Aneufinder, alterations were considered +1 when amplified, -1 when deleted, and 0 when unaltered. For the calculation of FGA, CNAs were considered to be the fraction of the genome for each arm based on its length in base pairs (extracted from reference genome hg38), positive when amplified and negative when deleted. FGA-G and FGA-L were derived by considering only positive or negative FGA values per sample, respectively.

### 5. Single-cell RNA sequencing

RPE-1 cells were plated at 30% confluency in 6 well plates (Greiner Bio-one) and treated for 16 hours with 62.5nM MPS1i followed by recovery for 32 hours. One million cells were harvested 48 hours after the start of treatment and frozen at -80 °C. Of note, these RPE-1 cells expressed a reporter of p53 activity, adapted from p53rep-iRFP-T2A-Luciferase plasmid (*61*) (gifted from the Karen H. Vousden lab). Cells were processed and sequenced by Single Cell Discoveries (Utrecht, The Netherlands). Briefly, sample preparation was performed in several steps according to an internal protocol: cryopreserved cells were thawed and then stained with AO/PI and evaluated on Luna Fx7 Automated cell counter. Viability was estimated to be at >90% for all of the samples in the group. Samples were further processed according to the Chromium GEM-X Single Cell 3’ v4 Gene Expression User Guide (CG000731). Libraries were sequenced on NovaSeq X Plus, 10B flowcell, 50-bp, paired end. 2 samples were sequenced: DMSO (5000 cells), MPS1i (20000 cells).

#### 5.1. Single-cell RNA sequencing analysis

Single-cell copy number analysis was performed in R (version 4.5.3) after processing, quality control, and filtering with Seurat (5.4.0) (*62*). For RPE-1 cells, DMSO- and Cpd5-treated cells were restricted to genes mapping uniquely to chromosomes 1-22 or X. Gene-level copy-number-associated expression shifts were estimated for each cell relative to clone-matched reference cells using DESeq2 (version 1.50.2) (*63*) negative-binomial models with size-factor normalization. Log2 fold changes were filtered by uncertainty, centered to the modal euploid signal within each cell, and segmented along chromosomes using Bayesian multiple-change-point models (mcp package, version 0.3.4) with clustermq parallelization (version 0.10.0) (*64*). Segment means were converted to integer copy-number states and summarized per cell as the number of altered chromosomes, the altered chromosome in single-event cells, and the genome-length-weighted fractions of gained or lost sequence. p53 pathway activity was inferred from log-normalized expression values using PROGENy (version 1.32.0) (*65*) with the top 500 pathway-responsive genes and normalized to mean 0/standard deviation 1. DMSO cells with detected aneuploidy were excluded. Genome-altered fractions were displayed for all eligible cells and separately for cells with at most one altered chromosome, using predefined bins for gained and lost genome fraction. Statistical significance was assessed using Student’s t test.

### 6. Visualization and statistical analysis

All plots and statistical analyses were performed in R (version 4.2.0) and R studio (version 23.12.1). For every statistical test, we assumed a significance threshold of 0.05 (corrected and uncorrected *p*-values). All R scripts used for data analysis in this study are available at: https://github.com/jrfmarques/TCGA_p53_aneuploidy.

#### 6.1. Modeling the relationship between p53 and FGA adjusted for confounders

To quantify the effects of p53 pathway deficiency on FGA or delta-FGA while accounting for clinical confounders, we used generalised linear models combined with model selection. We defined a hierarchical set of models including p53 pathway status, cancer stage, patient sex, patient age, and cancer subtype as covariates, together with all possible combinations of pairwise interaction terms between p53 pathway status and each covariate. This yielded 16 covariate-adjusted models, to which we added a p53-only model for comparison. Cancer stage, patient sex, and cancer subtype were treated as categorical variables.

A categorical variable was included for a given cancer type only when all represented categories had at least 10 observations; otherwise, the entire variable was excluded from modelling for that cancer type. Patient age was treated as a continuous variable and centered on the mean age within each cancer type. For the FGA analysis, we considered aneuploid samples without whole-genome doubling and excluded cancer types with fewer than 10 tumor samples. For delta-FGA, we considered cancer types with at least 10 tumor samples featuring both whole-chromosome and arm-level alterations (bCNAs).

Goodness-of-fit was assessed by fitting the simplest and most complex models in the pan-cancer dataset, excluding subtypes. For FGA, we evaluated Gamma and inverse Gaussian error distributions. For delta-FGA, we evaluated Gaussian, Student’s t, and fractional-logit models, and selected a Student’s t distribution as the best fit. Comparison of simulated and empirical residuals indicated that FGA was best described by a Gamma distribution with a log link including all main effects and interactions up to the five-way term, as defined in:

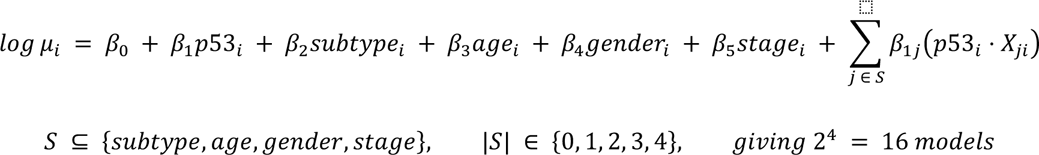

where FGA_i_ ∼ Gamma(μ_i_, ϕ), μ_i_ is the expected value of FGA for observation *i*, ϕ is the dispersion parameter, and S is the subset of covariates interacting with p53.

The 16 models defined by the choice of S:

**Model 1:** S = {} (no interactions; additive model)

**Model 2:** S = {subtype} → + β_1,sub_(p53 ⋅ subtype)

**Model 3:** S = {age} → + β_1,age_(p53 ⋅ age)

**Model 4:** S = {gender} → + β_1,gen_(p53 ⋅ gender)

**Model 5:** S = {stage} → + β_1,stg_(p53 ⋅ stage)

**Model 6:** S = {subtype, age} → + β_1,sub_(p53 ⋅ subtype) + β_1,age_(p53 ⋅ age)

**Model 7:** S = {subtype, gender} → + β_1,sub_(p53 ⋅ subtype) + β_1,gen_(p53 ⋅ gender)

**Model 8:** S = {subtype, stage} → + β_1,sub_(p53 ⋅ subtype) + β_1,stg_(p53 ⋅ stage)

**Model 9:** S = {age, gender} → + β_1,age_(p53 ⋅ age) + β_1,gen_(p53 ⋅ gender)

**Model 10:** S = {age, stage} → + β_1,age_(p53 ⋅ age) + β_1,stg_(p53 ⋅ stage)

**Model 11:** S = {gender, stage} → + β_1,gen_(p53 ⋅ gender) + β_1,stg_(p53 ⋅ stage)

**Model 12:** S = {subtype, age, gender} → + β_1,sub_(p53 ⋅ subtype) + β_1,age_(p53 ⋅ age) + β_1,gen_(p53 ⋅ gender)

**Model 13:** S = {subtype, age, stage} → + β_1,sub_(p53 ⋅ subtype) + β_1,age_(p53 ⋅ age) + β_1,stg_(p53 ⋅ stage)

**Model 14:** S = {subtype, gender, stage} → + β_1,sub_(p53 ⋅ subtype) + β_1,gen_(p53 ⋅ gender) + β_1,stg_(p53 ⋅ stage)

**Model 15:** S = {age, gender, stage} → + β_1,age_(p53 ⋅ age) + β_1,gen_(p53 ⋅ gender) + β_1,stg_(p53 ⋅ stage)

**Model 16:** S = {subtype, age, gender, stage} → + β_1,sub_(p53 ⋅ subtype) + β_1,age_(p53 ⋅ age) + β_1,gen_(p53 ⋅ gender) + β_1,stg_(p53 ⋅ stage)

For each cancer type, we fitted all 16 candidate models and computed the Akaike Information Criterion (AIC=−2ℓ(θ^) + 2k), where k includes all regression coefficients plus the Gamma dispersion parameter ϕ. The model with the lowest AIC among the covariate-adjusted models was selected for inference, while the p53-only model was retained for comparison but excluded from model selection (Supp. Table 5).

## Supplementary Materials

### Supplementary figures

**Supp. Fig. 1.**
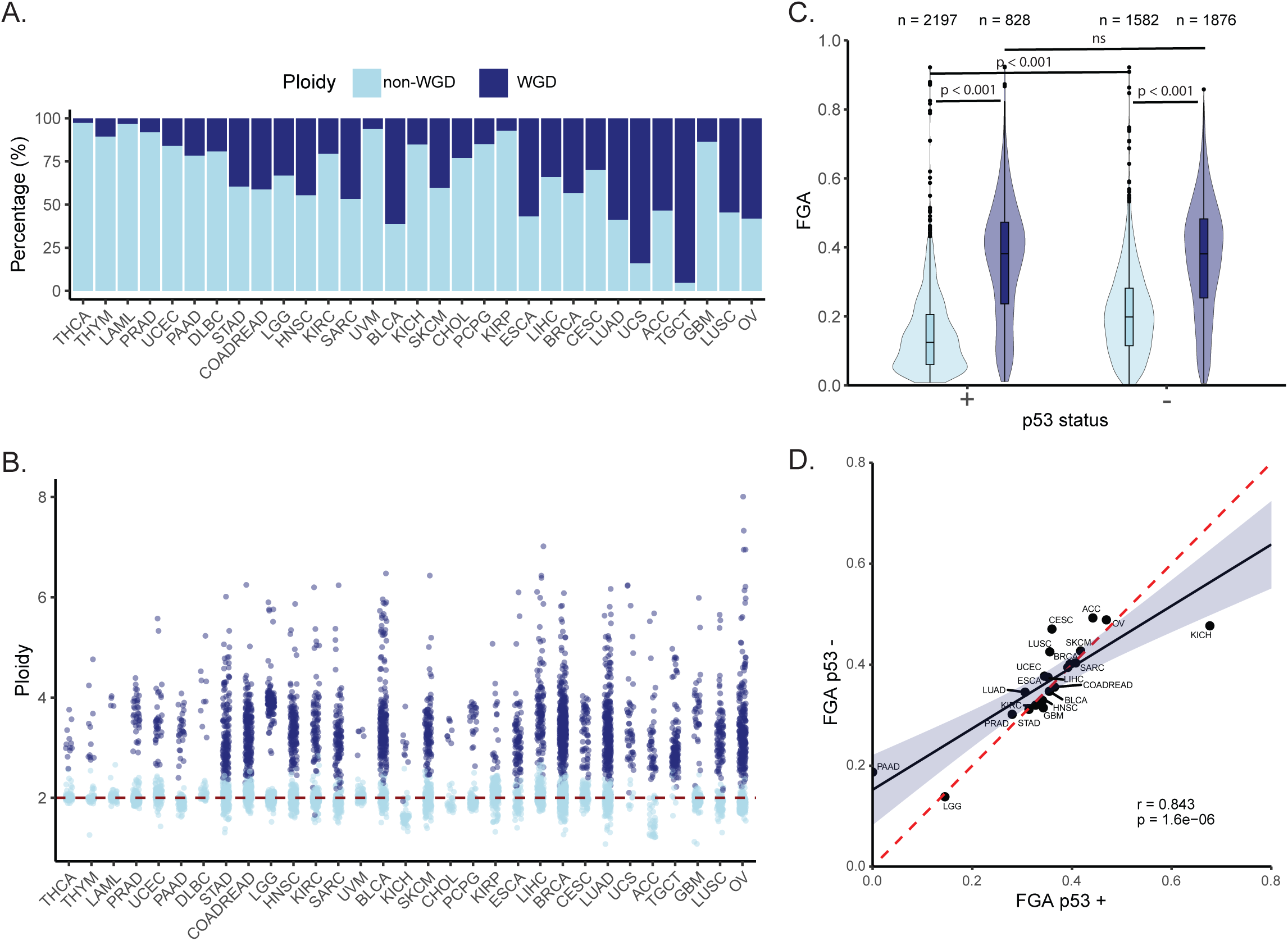
Distribution of WGD samples. A. Distribution of WGD (dark blue) and non-WGD events (light blue) per cancer type. Tumors are ordered based on average aneuploidy frequency. B. Distribution of ploidy per cancer type. Colors depict samples with WGD (dark blue) and non-WGD event (light blue). Tumors are ordered based on average aneuploidy frequency. Doted red line marks diploidy (2N). C. Pan-cancer FGA distribution of samples with WGD (dark blue) and non-WGD events (light blue) divided into p53 proficient (+) and deficient (-) samples. Boxplot represents the median and first and third quartiles. Dots represent outliers. “n=” number of tumor samples per group. D. Distribution of FGA in Tp53 proficient (+) and deficient (-) tumors, per cancer type in samples with WGD event. Linear regression line (black) and confidence interval (dark blue shade). Doted red line depicts a 1:1 ratio. Pearson’s correlation test was used.

**Supp. Fig. 2.**
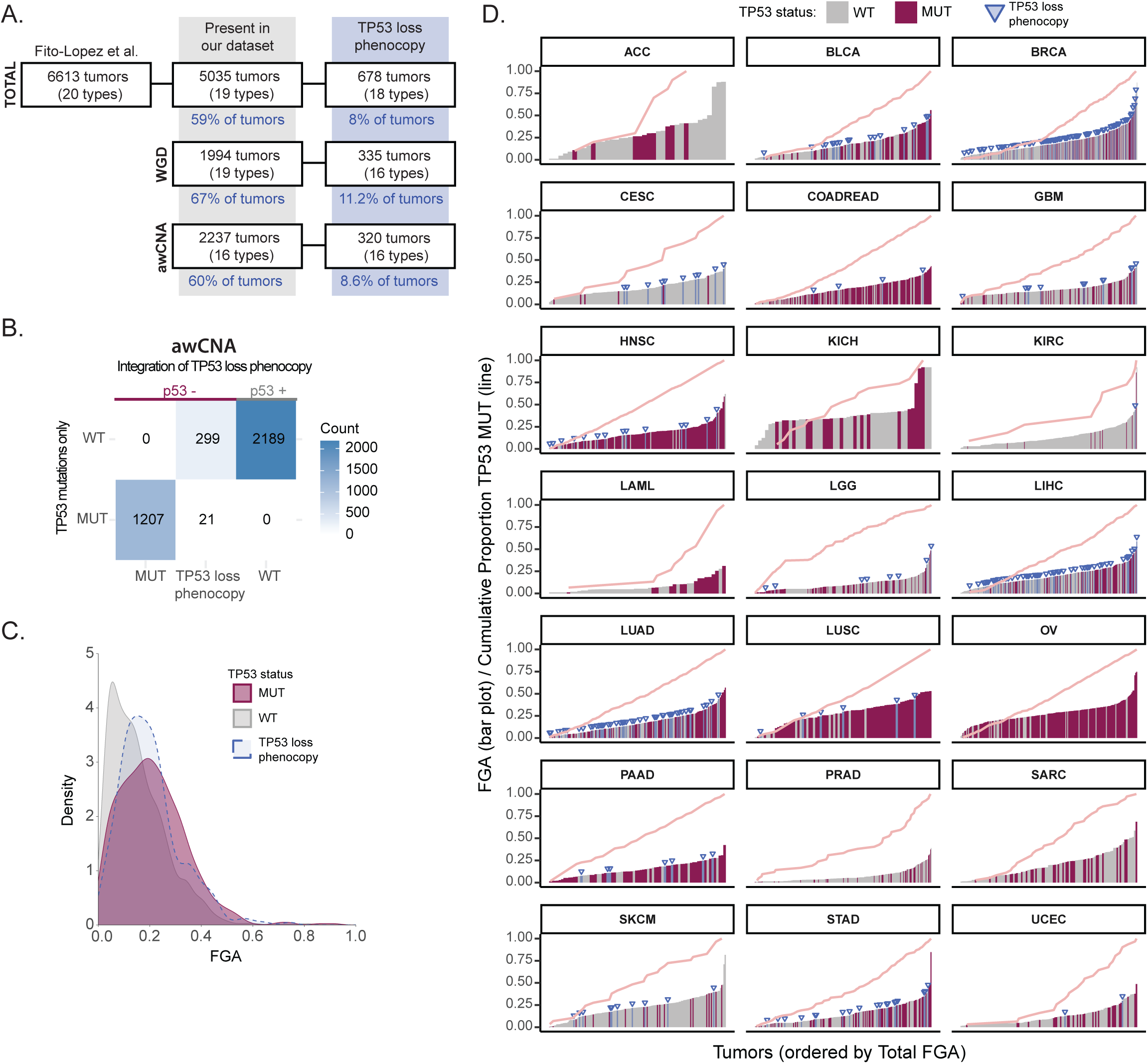
Description of samples with TP53 loss phenotype. A. Number of samples from dataset published Fito-Lopez e al. that match our current dataset and are classified as “TP53 loss phenocopy” across WGD and awCNA tumors. B. Distribution and integration of “TP53 loss phenocopy” samples as p53 deficient samples in our dataset. Concordance table of classification of samples based on TP53 mutation status (MUT or WT) and samples defined as TP53 loss phenocopy. On top shows the final TP53 status grouping used for analysis where p53 deficient (-) includes TP53 mutated and TP53 loss phenocopy tumors and p53 proficient (+) includes only TP53 WT tumors. C. Pan-cancer FGA distribution of TP53 WT (grey) MUT (pink) for awCNA tumors according to mutation status only. TP53 loss phenocopy are highlighted in blue. Density histogram in arbitrary units (AU), AUC=1 for each variable. D. Distribution of tumor samples in ascending order of FGA for each cancer type. The color filling corresponds to TP53 status – WT (grey) and MUT (pink). TP53-loss phenocopy tumors are highlighted in blue with a blue arrow head. Line graphs (light pink) show the relative cumulative frequency of TP53 mutated samples.

**Supp. Fig. 3.**
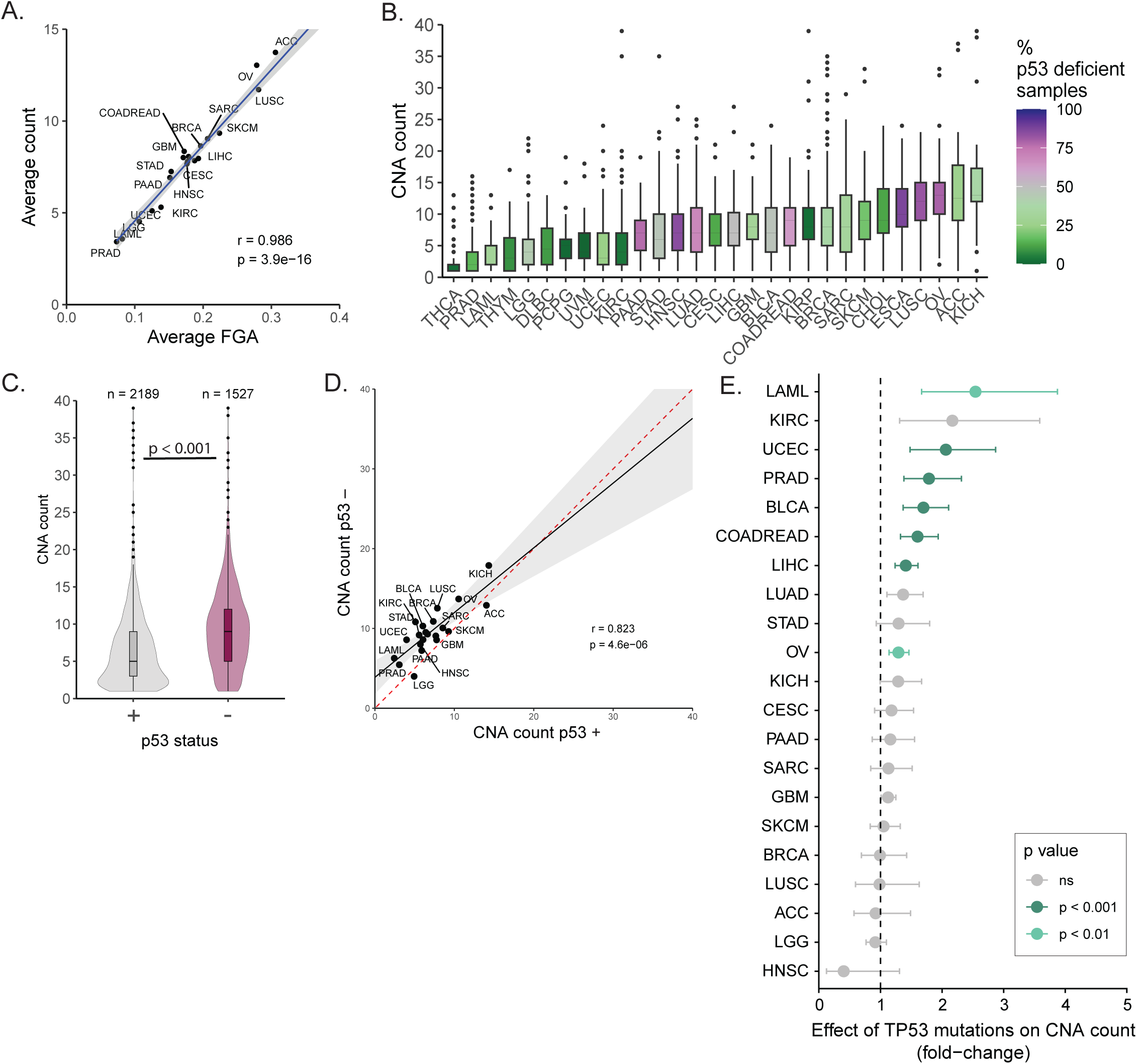
Distribution of copy number alteration counts for awCNA. A. Distribution of average FGA and average CNA count (per chromosome arm) in bCNA samples, per cancer type. Linear regression line (blue) and confidence interval (grey shade). Pearson’s correlation test was used. B. CNA counts per arm for awCNA events per cancer type (ordered by mean CNA count). Color filling depicts the % of p53 deficient samples per cancer type. C. Pan-cancer CNA counts distribution in p53 proficient (+, grey) and deficient (-, pink). Boxplot represents the median and first and third quartiles. Dots represent outliers. “n=” number of tumor samples per group. Statistical analysis was performed using Mann-Whitney-U test. D. Distribution of CNA counts in p53 proficient (+) and deficient (-) tumors, per cancer type. Linear regression line (black) and confidence interval (grey shade). The doted red line depicts a 1:1 ratio. Pearson’s correlation test was used. F. Fold-change of effect size of p53 deficiency on CNA counts of tumors, per cancer type. Estimated using a generalized linear model (GLM) with gamma distribution. Colors indicate the significance of the effect: bars are colored by p-value threshold (p<0.05, p<0.01, p<0.001) and shown in grey when the effect is not significant (ns). Error bars represent the standard error of the model coefficient, accounting for overdispersion in the Gamma GLM framework.

**Supp. Fig. 4.**
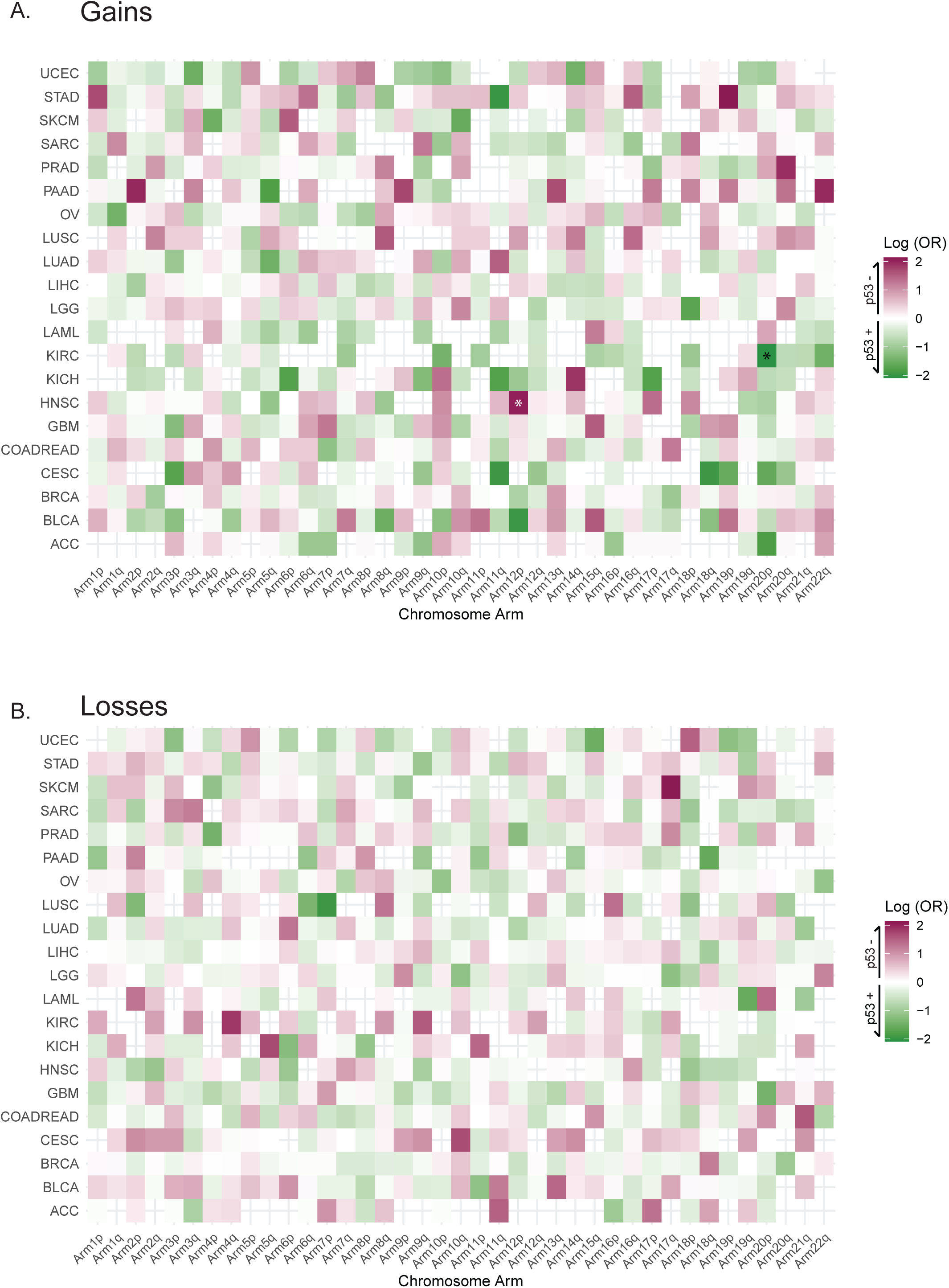
Distribution of FGA attributed to gains and losses per cancer type and chromosome arm in relation to p53 status. A. Correlation of FGA attributed to gains and p53 deficiency per cancer type and chromosome arm, for awCNAs. Color fill shows log(odds ratio) of gains versus no gains: positive values (pink gradient) depict FGA enriched in p53 deficient samples and negative values (green gradient) depicts FGA enriched in p53 proficient samples. Analysis was performed using Fisher’s exact test. Bonferroni correction for multiple testing was applied across all arms for each cancer type. Asterisk (*) represents significant arms. B. Correlation of FGA attributed to losses and p53 deficiency per cancer type and chromosome arm, for awCNAs. Color fill shows log(odds ratio) losses vs no losses: positive values (pink gradient) depict FGA enriched in p53 deficient samples and negative values (green gradient) depicts FGA enriched in p53 proficient samples. Analysis was performed using Fisher’s exact test. Bonferroni correction for multiple testing was applied across all arms for each cancer type. No significant correlations were found.

**Supp. Fig. 5.**
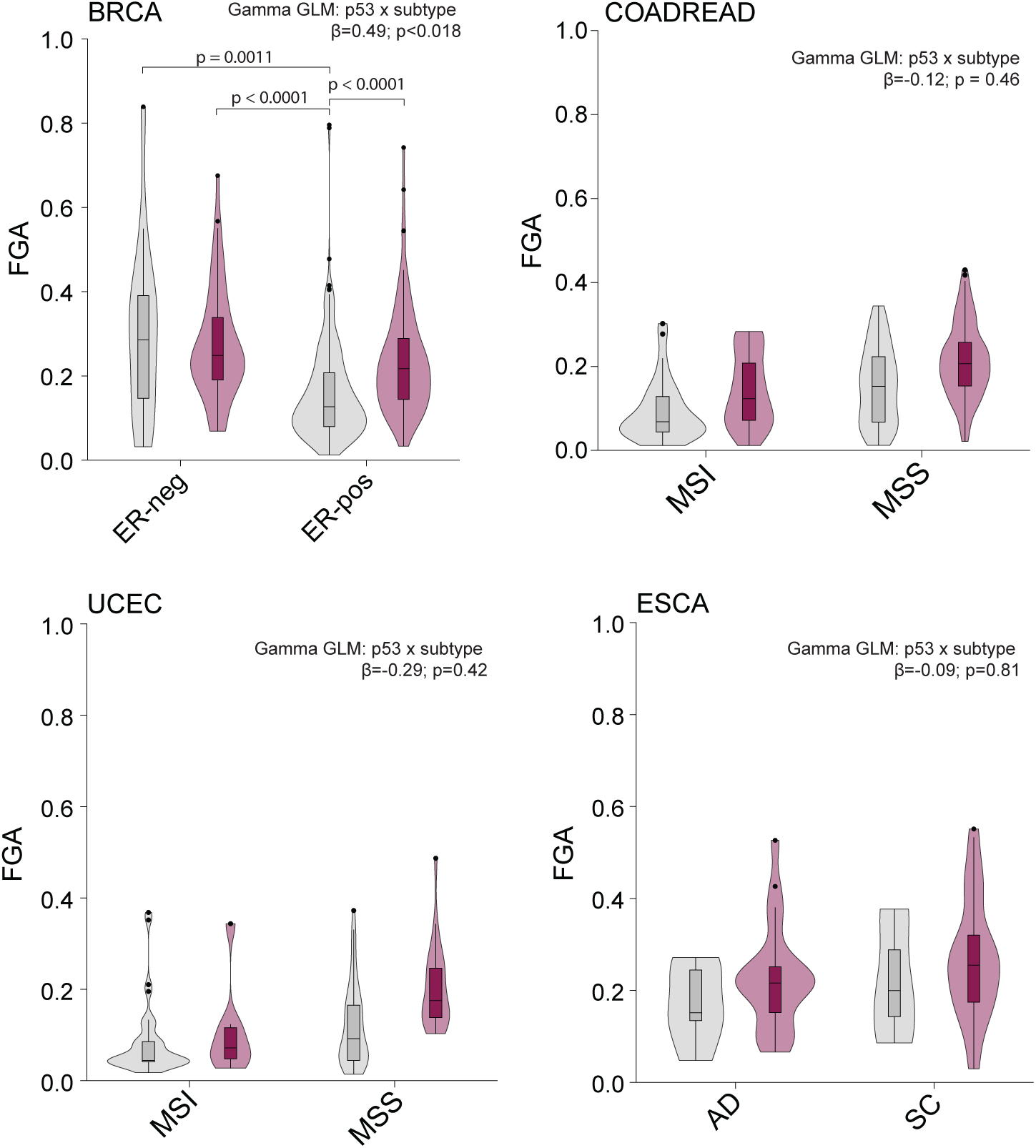
Effect of subtype and p53 on FGA distribution. Distribution of FGA in p53-deficient (pink) and proficient (grey) for cancer subtypes of BRCA: estrogen receptor positive (ER-pos) and negative (ER-neg); COADREAD: Microsatellite Instability (MSI) and Microsatellite Stable (MSS); UCEC: Microsatellite Instability (MSI) and Microsatellite Stable (MSS); ESCA: Adenocarcinoma (AD) and Squamous cell (SC). Gamma GLM summary results for the interaction of p53 status and subtype are shown in the top right corner of each plot. For BRCA, the only cancer type with significant interaction, significant pairwise comparisions are shown.

**Supp. Fig 6.**
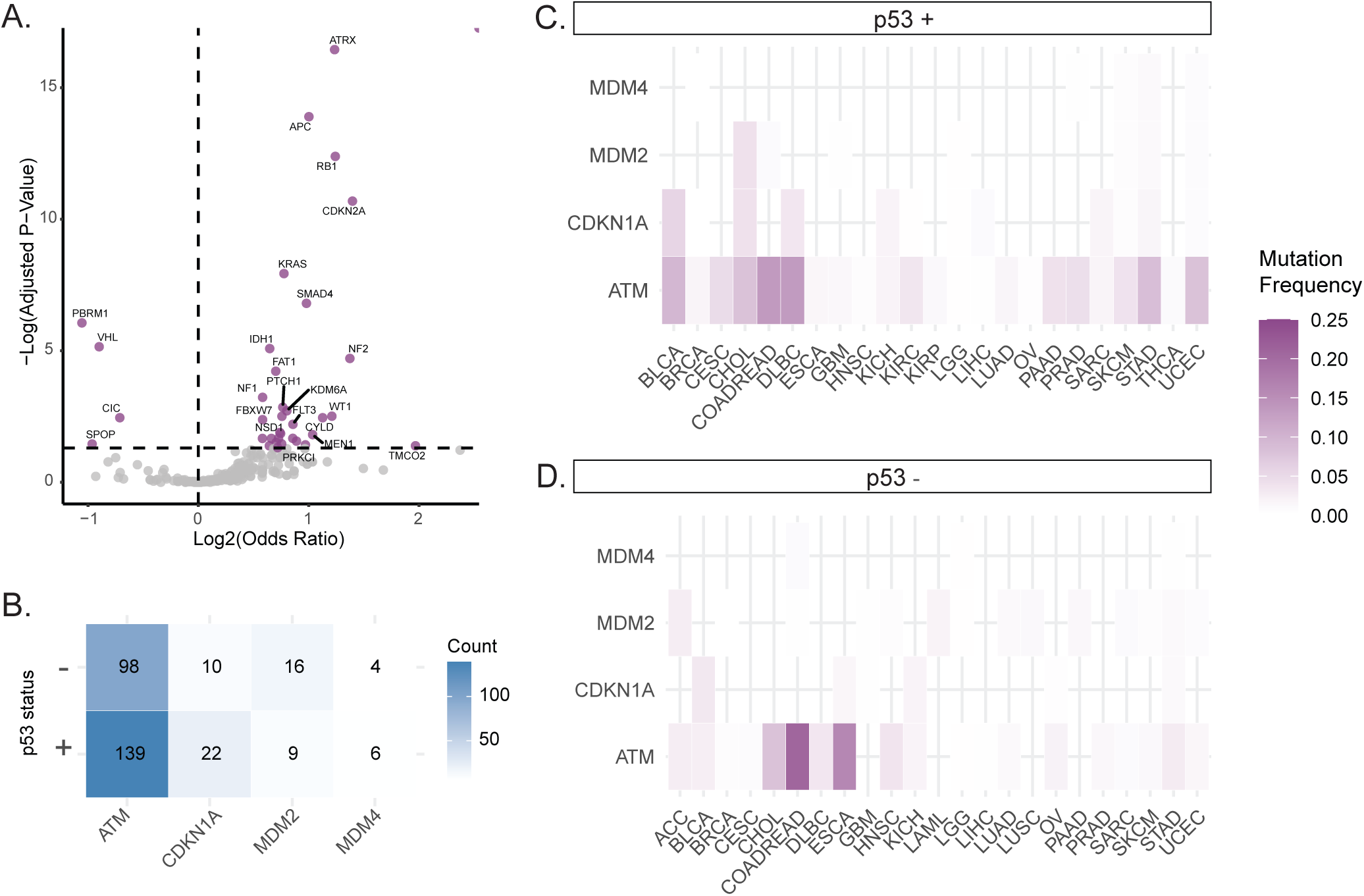
Prevalence of mutations in p53 pathway and correlation between TP53 mutations and other driver mutations. A. Association of p53 deficiency and most significant driver genes in cancer (see methods). Statistical analysis was performed using Fisher’s exact test. Calculated p-values were corrected for multiple testing using the Bonferroni method. Genes in purple are significant correlated (positive odds ratio) or anti-correlated (negative odds ratio) with occurrence of TP53 mutations. B. Distribution of samples with mutations in genes associated to the TP53 pathway divided by p53 proficient and deficient awCNA samples. Color gradient represents the number of samples. C. Distribution of p53 proficient samples mutated for TP53 pathway associated genes, per cancer type. Color gradient represents frequency of mutated samples for a given gene. D. Distribution of p53 deficient samples mutated for TP53 pathway associated genes, per cancer type. Color gradient represents frequency of mutated samples for a given gene.

**Supp. Fig. 7.**
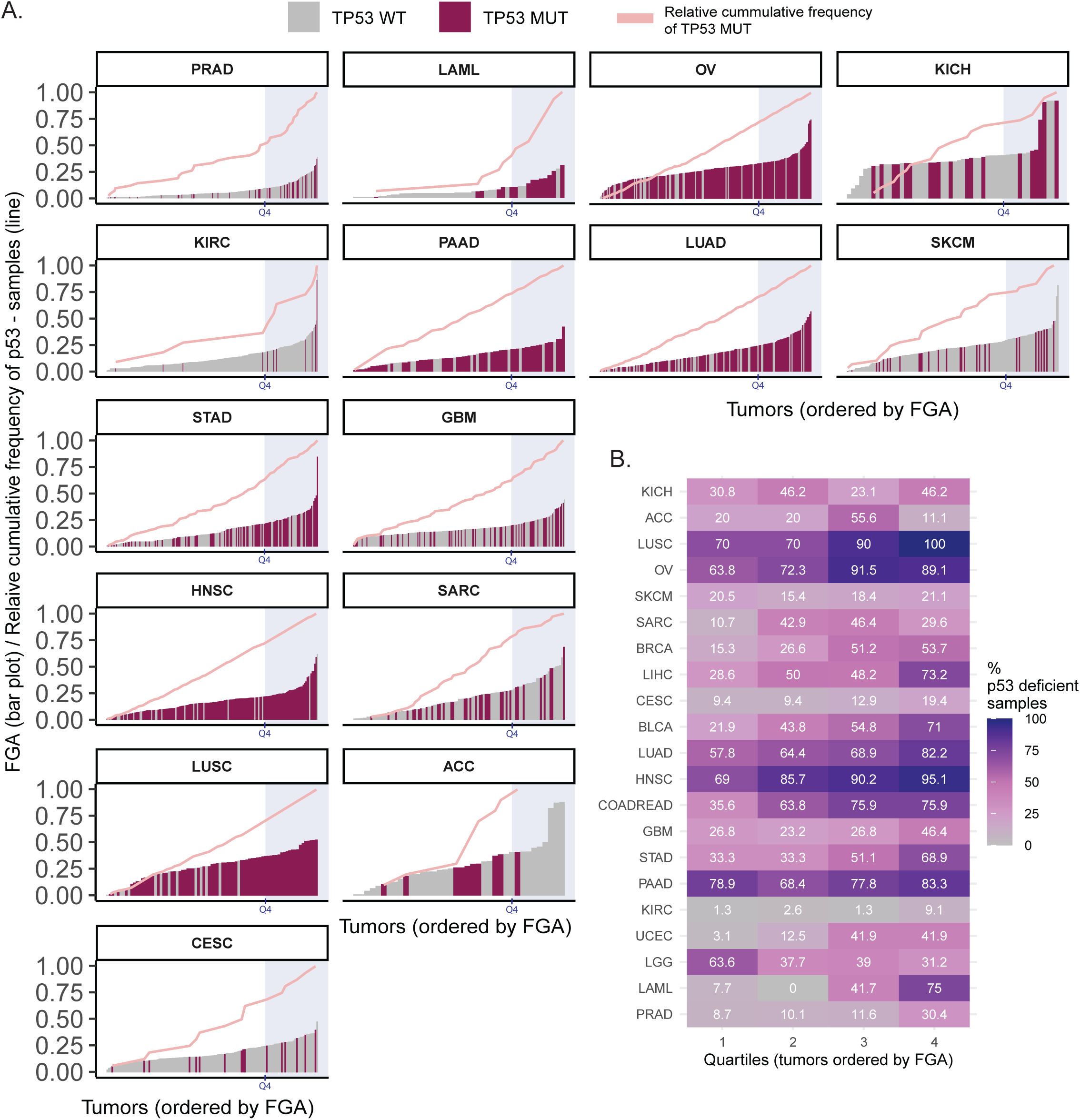
Quantification of p53 deficiency distribution in tumors of each cancer type. A. Distribution of tumor samples in ascending order of FGA for each cancer type. The color filling corresponds to p53 status – proficient (grey) and deficient (pink). Line graphs (light pink) show the relative cumulative frequency of p53 deficient (-) samples. Shaded regions (light blue) correspond to the fourth quartile for each cancer type. B. Distribution of p53 deficient samples grouped by quartiles of samples in ascending order of FGA, for each cancer type. Color filling corresponds to the percentage of p53 deficient samples for each quartile.

**Suppl. Fig. 8.**
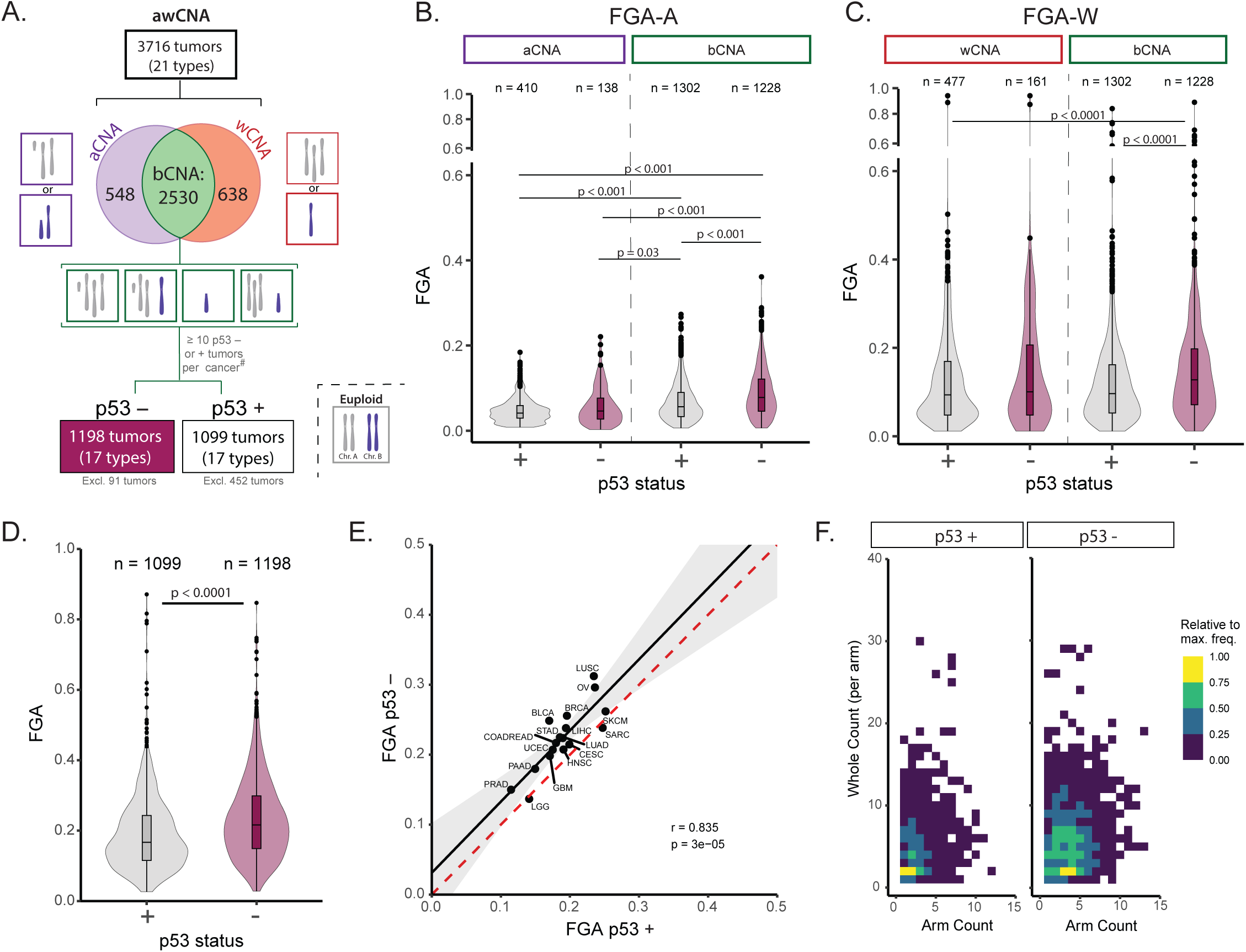
p53 deficiency associates neither with arm-level nor whole chromosome CNAs. A. awCNA were subcategorized based on the type of aneuploidy into samples with arm-level only (aCNA), whole chromosome only (wCNA) and both alterrations (bCNA). The euploid karyotype depicts 2 chromosome pairs: the grey pair is used to depict whole-chromosome or arm gains and the blue pair represents losses. For each category example of karyotypes are shown. bCNA samples were further filtered based on p53 status. Grey text shows excluded samples for each category.TP53 B. Pan-cancer FGA attributed to arm-level alterations (FGA-A) in samples with only arm-level alterations (aCNA) or in combination with whole chromosome alterations (bCNA), in p53 proficiency (grey) and deficiency (pink). Boxplot represents the median and first and third quartiles. Dots represent outliers. “n=” number of tumor samples per group. Statistical analysis was performed using pairwise Mann-Whitney-U test (Bonferroni correction method). C. Pan-cancer FGA attributed to whole chromosome alterations (FGA-W) in samples with only whole chromosome alterations (aCNA) or in combination with arm-level alterations (bCNA), in p53 proficiency (grey) and deficiency (pink). Boxplot represents the median and first and third quartiles. Dots represent outliers. “n=” number of tumor samples per group. Statistical analysis was performed using pairwise Mann-Whitney-U test (Bonferroni correction method). D. Distribution of FGA in p53 proficiency and p53 deficiency in bCNA tumors per cancer type. Linear regression line (black) and confidence interval (grey shade). Doted red line depicts a 1:1 ratio. Pearson’s correlation test was used. E. Distribution of FGA in p53-proficient and p53-deficient bCNA tumors, per cancer type. Linear regression line (black) and confidence interval (grey shade). The doted red line depicts a 1:1 ratio. Pearson’s correlation test was used. F. Frequency distribution of whole-chromosome and am-level event counts relative to the most frequent event, in p53-proficient samples (left) and p53-deficient samples (right). Whole chromosome events are depicted as 2 arm-level events.

**Supp. Fig. 9.**
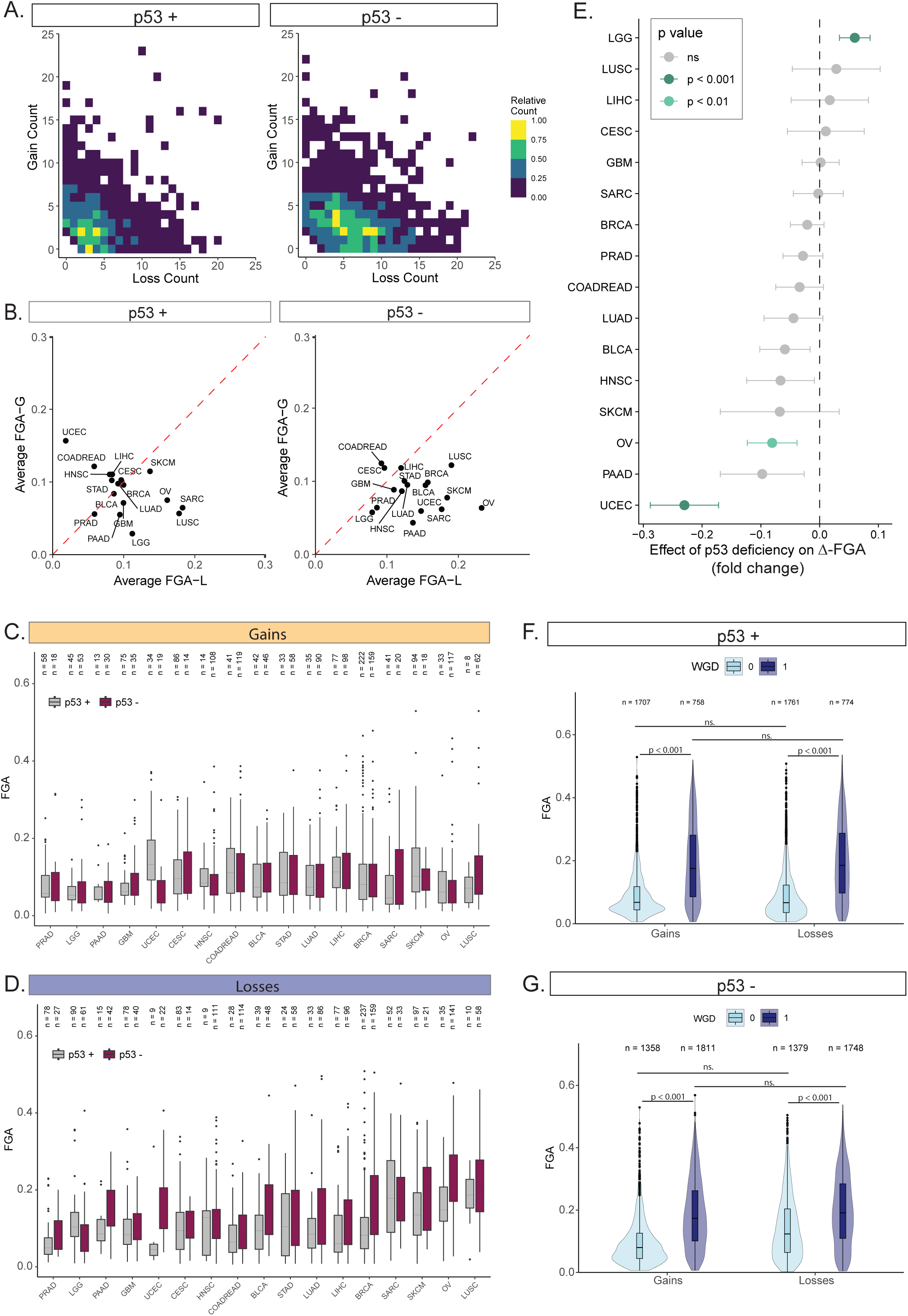
Distribution of copy-number gains and losses. A. Frequency distribution of gain and loss event counts relative to the most frequent event, in p53 proficient samples. Whole chromosome events are depicted as 2 arm-level events. B. Left: Distribution of FGA-G vs FGA-L, per cancer type, in p53-proficient samples. Doted red line depicts a 1:1 ratio. Right: Distribution of FGA-G vs FGA-L, per cancer type, in p53-deficient samples. Doted red line depicts a 1:1 ratio. C. Distribution of FGA-G in p53 proficient and deficient samples per cancer type. “n=” number of tumor samples per group. D. Distribution of FGA-L in p53 proficient and deficient samples per cancer type. “n=” number of tumor samples per group. E. Fold-change of effect size of p53 deficiency on FGA of tumors, per cancer type. Estimated using a generalized linear model (GLM) with gamma distribution. Colors indicate the significance of the effect: bars are colored by p-value threshold (p<0.05, p<0.01, p<0.001) and shown in grey when the effect is not significant (ns). Error bars represent the standard error of the model coefficient, accounting for overdispersion in the Gamma GLM framework. F. Pan-cancer FGA-G and FGA-L of p53 proficient samples with WGD (dark blue) and non-WGD events (light blue). Boxplot represents the median and first and third quartiles. Dots represent outliers. “n=” number of tumor samples per group. G. Pan-cancer FGA-G and FGA-L of p53 deficient samples with WGD (dark blue) and non-WGD events (light blue). Boxplot represents the median and first and third quartiles. Dots represent outliers. “n=” number of tumor samples per group.

**Supp. Fig. 10.**
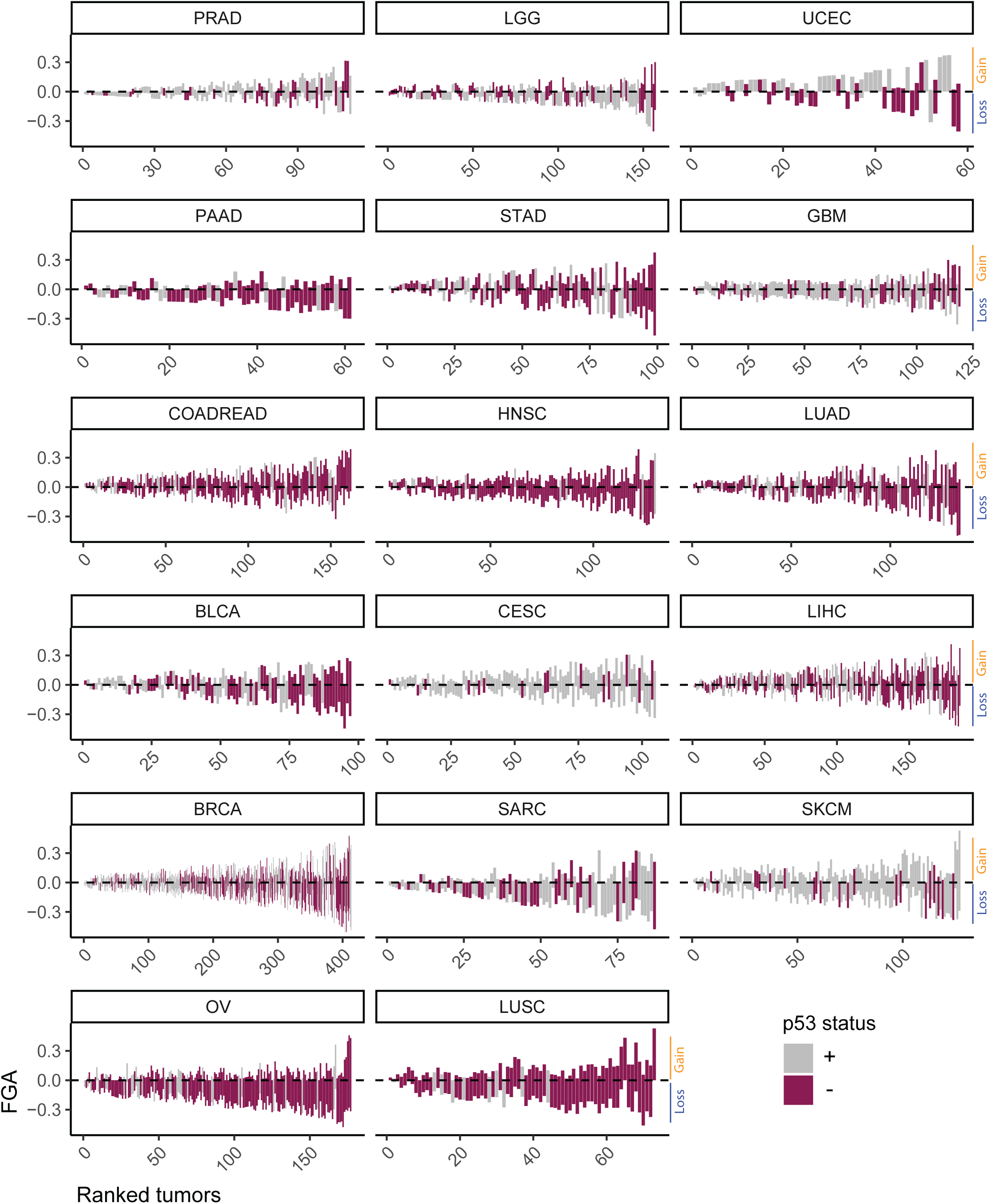
Distribution of FGA divided into gains and losses in bCNA samples per cancer type. Distribution of FGA-G (positive FGA) and FGA-L (negative FGA) per tumor sample across cancer types. Samples are distributed in ascending order of total FGA. The color filling corresponds to p53 status – proficient (+, grey) and deficient (-, pink).

**Supp. Fig. 11.**
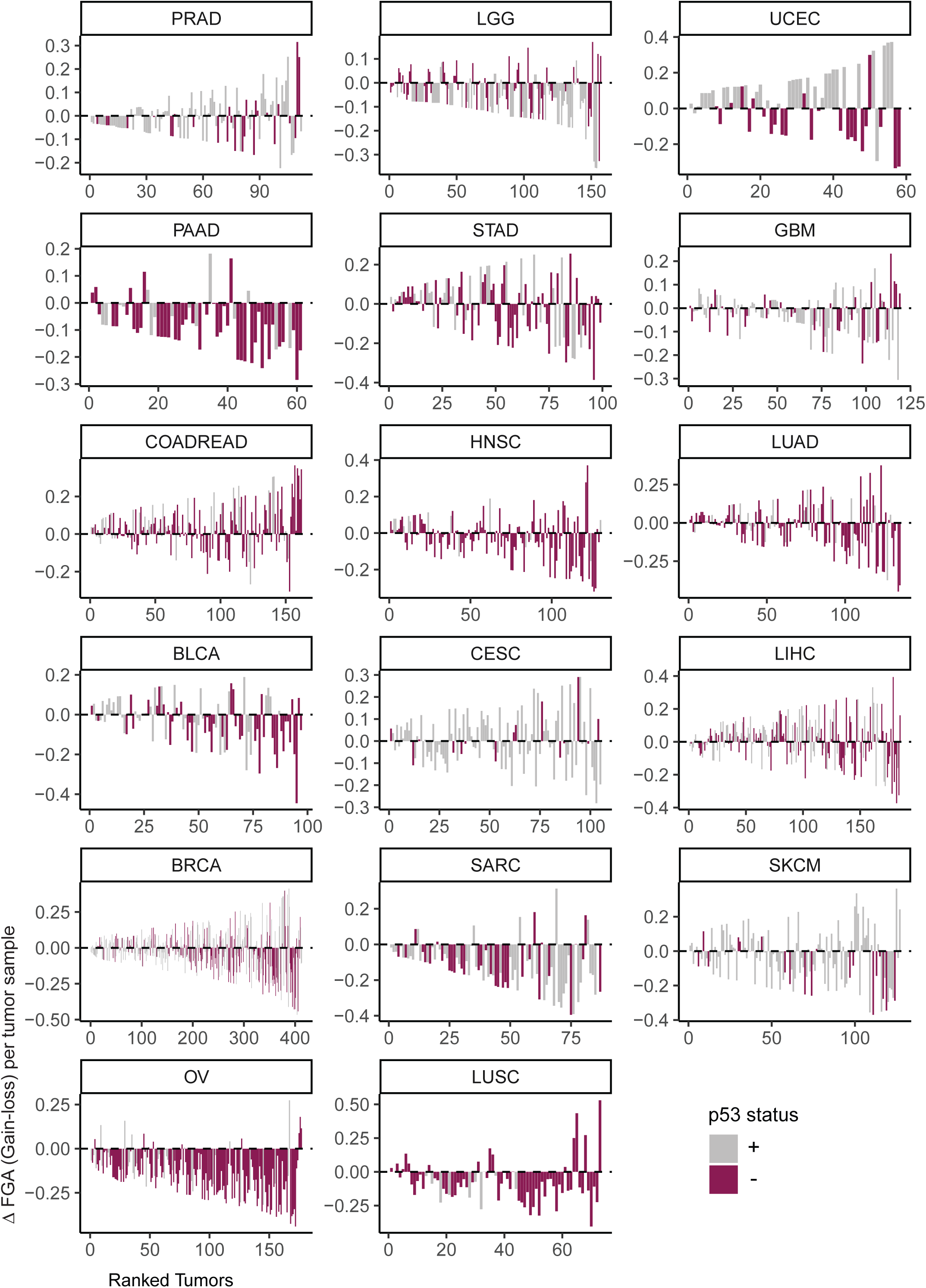
Delta FGA (FGA-G minus FGA-L) in bCNA tumors per cancer type. Delta of FGA (FGA-G minus FGA-L) per tumor sample across cancer types. Samples are distributed in ascending order of total FGA. The color filling corresponds to p53 status – proficient (grey) and deficient (pink). Positive values correspond to the higher burden of FGA-G and negative values to a higher burden of FGA-L in each tumor sample.

**Suppl. Fig. 12.**
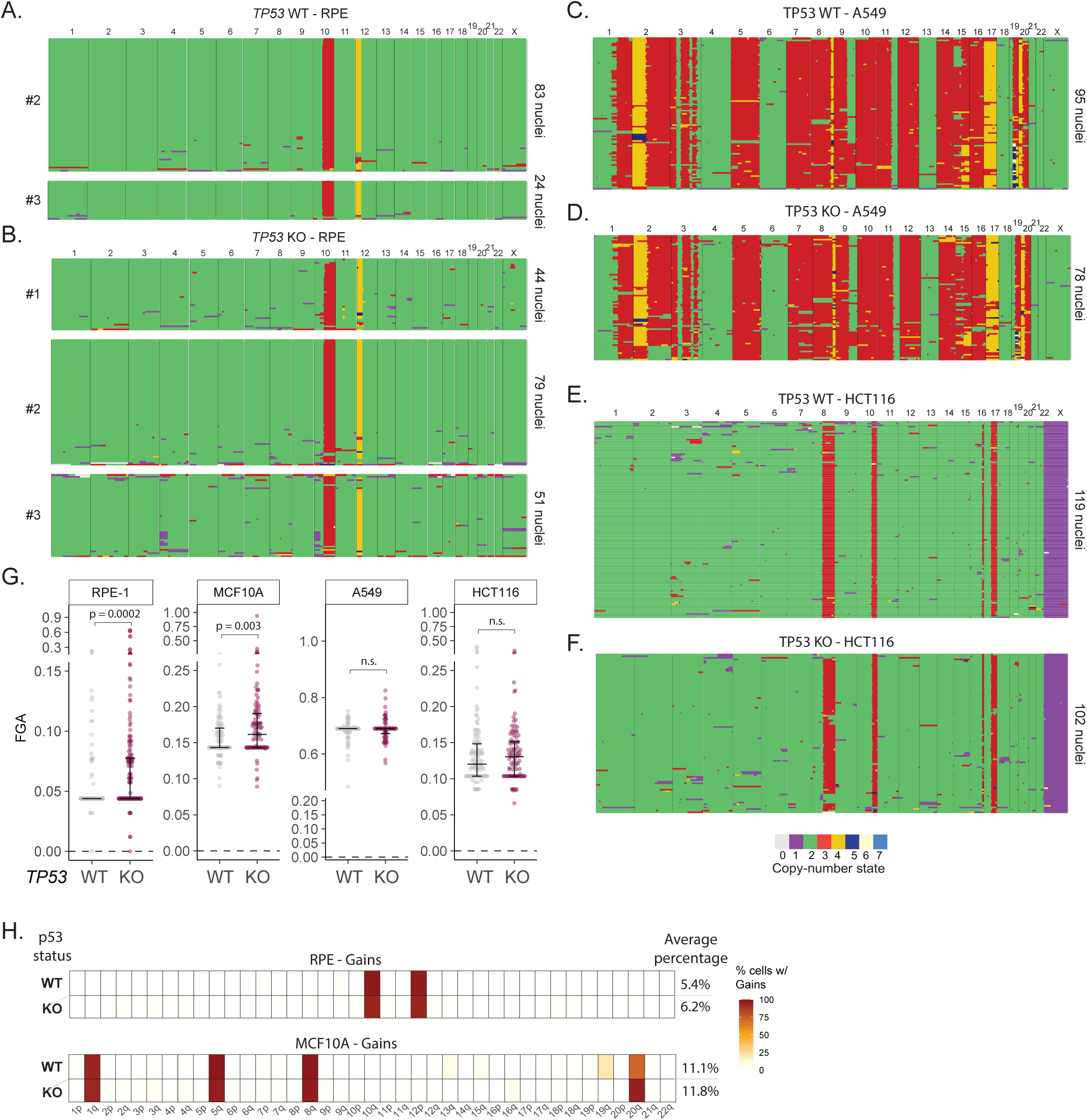
Distribution of FGA in non-transformed and transformed cell lines. (A-B) Single cell karyotype sequencing plots for RPE-1 cell line. Conditions: TP53 WT (A) and TP53 KO (B). Color code depicts copy number states. 2-3 replicates were used per condition represented by #. Right: Number of sequenced nuclei. (C-D) Single cell karyotype sequencing plots for A549 cell line. Conditions: TP53 WT (C) and TP53 KO (D). Color code depicts copy number states. 1 replicate per condition. Right: Number of sequenced nuclei. (E-F) Single cell karyotype sequencing plots for HCT116 cell line. Conditions: TP53 WT (E) and TP53 KO (F). Color code depicts copy number states. 1 replicate per condition. Right: Number of sequenced nuclei. (G) Distribution of FGA across cell lines ((RPE, MCF10A, A549 and HCT116) with TP53 WT (grey) or KO (pink). Data corresponds to 2 to 3 replicates for RPE-1 cells and one replicate for the othr lines. Kruskal-Wallis test with pairwise Mann-Whitney U tests and Bonferroni correction for multiple comparisons was used. (H) Average frequency of gain events per chromosome arm in RPE-1(top) and MCF10A (bottom) cells with TP53 WT or KO. Right: average percentage of cells with gain events per condition.

**Suppl. Fig. 13.**
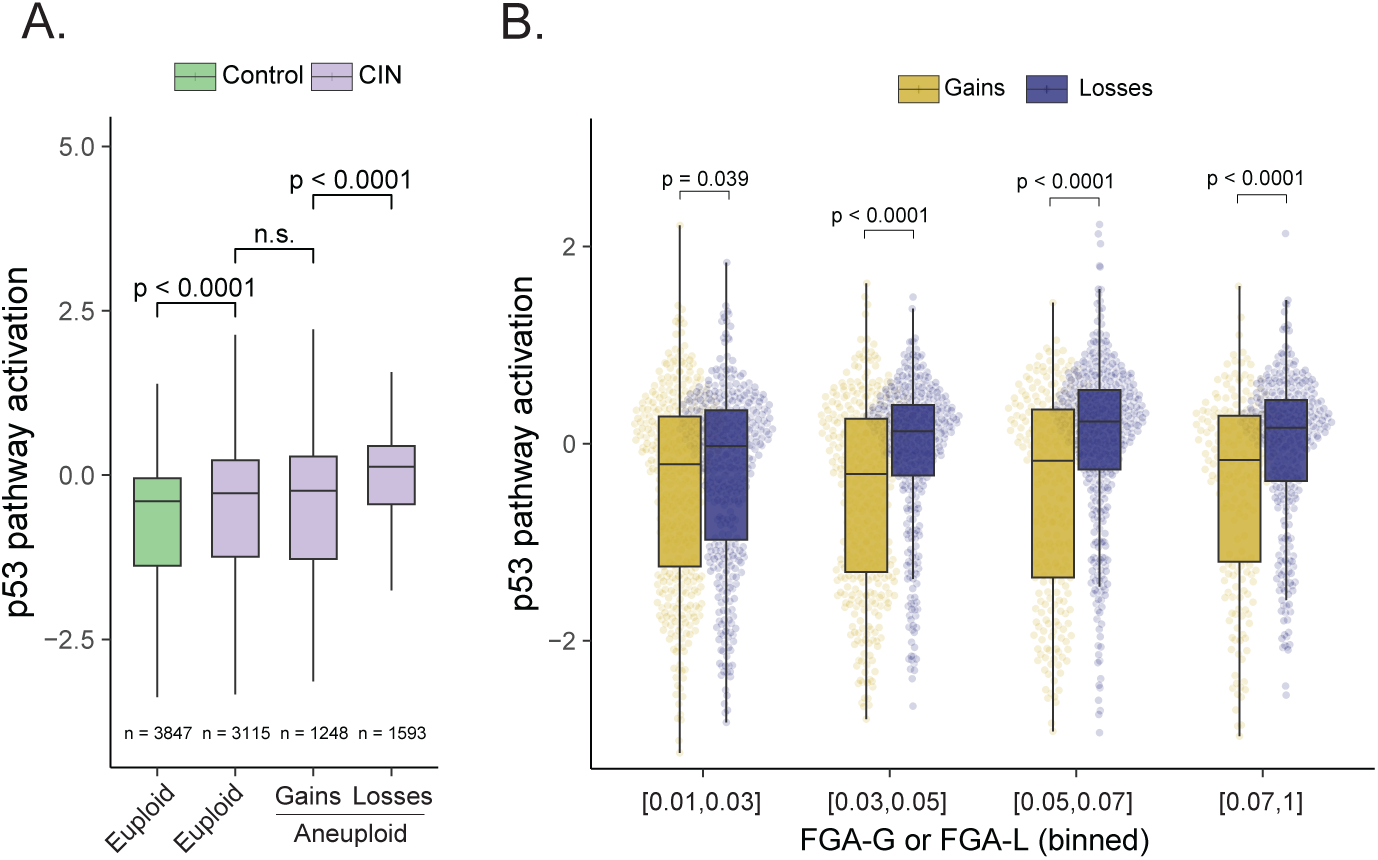
p53 response to single whole-chromosome gains and losses. (A) Box-plot of p53 pathway activation score for euploid (FGA = 0) control (green) and euploid, whole-chromosome gains (one event per cell at most) and whole-chromosome losses only (one event per cell at most) CIN-induced cells (purple). Box plot represents median and first and fourth quartile. n = the number of cells per condition. Student’s t test was used. (B) p53 pathway activation score for cells with at most one whole-chromosome gain (yellow) or one whole-chromosome loss per cell (blue), binned per increasing degree of FGA. Each dot represents one single cell. Box plot represents median and first and fourth quartile. Student’s t test was used.

**Suppl. Fig. 14.**
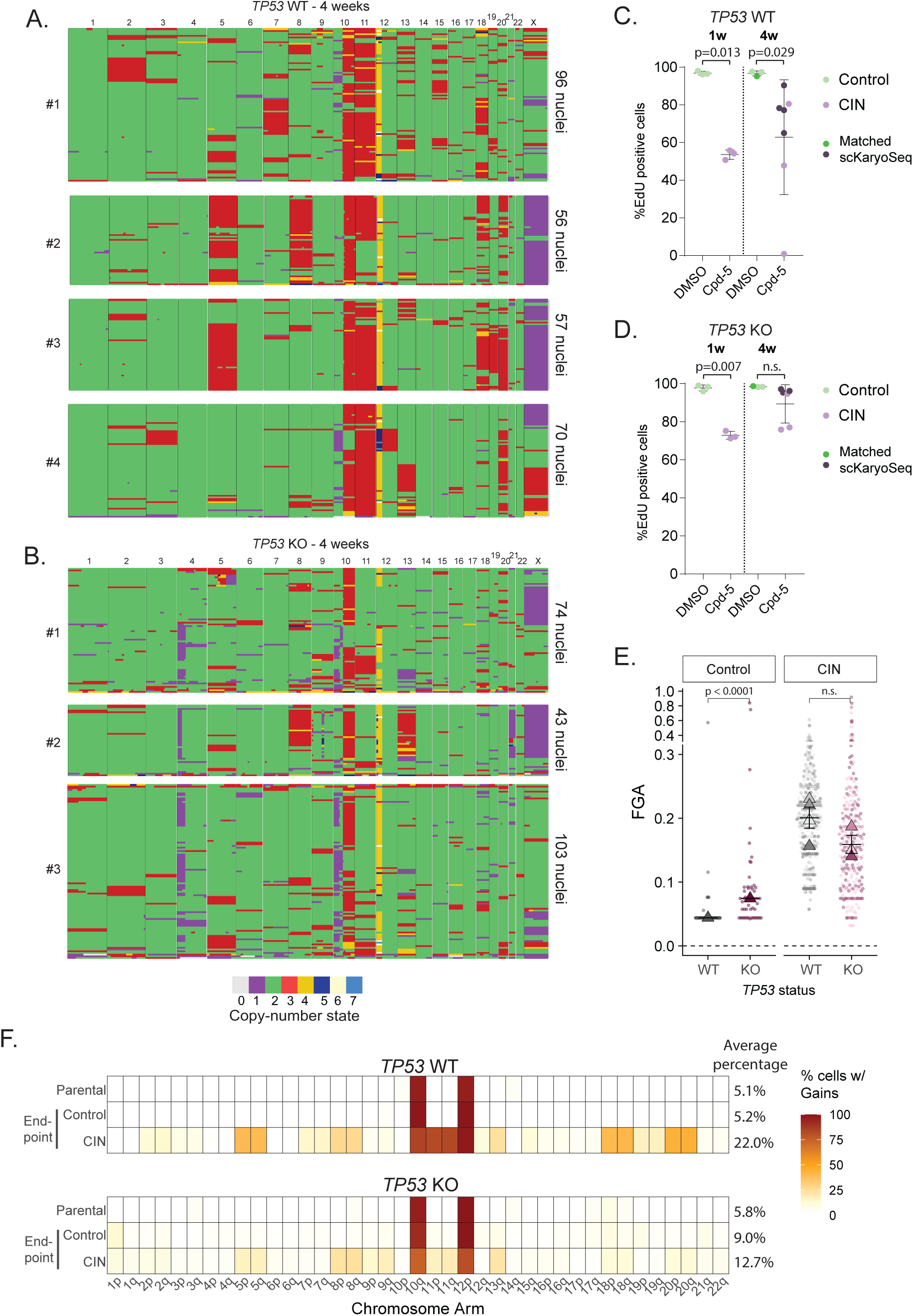
Effect of p53 inactivation in long-term evolution of RPE-1 cells. (A-B) Single cell karyotype sequencing plots for RPE-1 cell line treated with Cpd-5 for 4 weeks. Conditions: *TP53* WT (A) and *TP53* KO (B). Color code depicts copy number states. 3-4 replicates were used per condition represented by #. Right: Number of sequenced nuclei. (C-D) Proliferation rate (measured as percentage of EdU positive cells after 24h pulse) of *TP53* WT ((C) and *TP53* KO (D) cells at 1 week or 4 week timepoint after DMSO (Control - green) and Cpd-5 (CIN - purple) treatment. Darker colors depict samples with matched scKaryoSeq data in (A-B) and Fig. 5D,E. Kruskal-Wallis test with pairwise Mann-Whitney U tests and Bonferroni correction for multiple comparisons was used. (E) Distribution of FGA in RPE with TP53 WT (grey) or KO (pink), for control and CIN conditions at endpoint (4 weeks). Each dot represents an individual cell, coloured by replicate, when applicable. *TP53* WT (shades of grey) and *TP53* KO (shades of pink).Triangles indicate the median value per replicate. Horizontal bars represent the mean of replicate medians ± standard error of the mean (SEM). Control has only one replicate; the bar represents the median of all cells ± SEM. Statistical comparisons between SCR and KO within each condition were performed using a two-sided Mann-Whitney U test. For control condition, all individual cells were used as observations. For CIN condition, replicate medians were used with each replicate representing an independent experiment. (F) Average frequency of gains per chromosome arm in RPE-1 cells with *TP53* WT or KO, for parental and endpoint conditions (control and CIN treated). Right: Average percentage of cells with gain events per condition.

### Supplementary Tables

**Supp. Table 1. Overview of number of samples per cancer type.**

A. Cancers name abbreviation as defined in TCGA. Summary of number of euploid, aneuploid and p53 deficient samples and respectives percentages, per cancer type.

B-F. Summary of number of aneuploid and p53 deficient samples for each category: WGD (B), awCNA (C), aCNA (D), wCNA (E), bCNA (F).

**Supp. Table 2. Compilation of copy-number counts and FGA per tumor sample.**

A-E. Copy-number counts and FGA (sum) per tumor sample for all samples unfiltered (A) (including WGD+ samples), for awCNA (B), aCNA (C), wCNA (D), bCNA (E).

**Suppl. Table 3. Average number distribution of CNAs per p53 status.**

Average copy number counts and FGA (sum) grouped by cancer type and p53 status for WGD (A), aCNA (B), wCNA (C), and bCNA (D).

**Suppl. Table 4 - Statistical tests results.**

A. Pairwise fisher exact test applied in fig. 1C. Results show p value, odds ratio and adjusted p value (using bonferroni correction) per cancer type.

B. Pairwise fisher exact test applied in fig. 1E. Results show p value, odds ratio and adjusted p value (using bonferroni correction) per cancer type.

C. Pairwise Mann-Whitney U test applied in fig. 2G. Results show p value, odds ratio and adjusted p value (using bonferroni correction) per cancer type and quartile (Q2-4) relative to quartile 1.

D. Pairwise Mann-Whitney U test applied in fig. 3B. Results show p value and adjusted p value (using bonferroni correction) per cancer type.

E. Pairwise Mann-Whitney U test applied in fig. 3C. Results show p value and adjusted p value (using bonferroni correction) per cancer type.

F. Pairwise Mann-Whitney U test applied in Suppl. fig. 1C. Results show p value for each comparison.

**Suppl. Table 5 - Selected model results for p53 effect on FGA adjusted for confounders per cancer type.**

Selected model results for p53 effect on FGA corrected for subtype, stage, age and sex as confounders, per cancer type. TRUE/FALSE matrix depicts the confounders assessed per model and cancer type. Stage was considered as a factor.

